# Nucleotide-resolution bacterial pan-genomics with reference graphs

**DOI:** 10.1101/2020.11.12.380378

**Authors:** Rachel M Colquhoun, Michael B Hall, Leandro Lima, Leah W Roberts, Kerri M Malone, Martin Hunt, Brice Letcher, Jane Hawkey, Sophie George, Louise Pankhurst, Zamin Iqbal

## Abstract

**Background:** Bacterial genomes follow a U-shaped frequency distribution whereby most genomic loci are either rare (accessory) or common (core); the union of these is the pan-genome. The alignable fraction of two genomes from a single species can be low (e.g. 50-70%), such that no single reference genome can access all single nucleotide polymorphisms (SNPs). The pragmatic solution is to choose a close reference, and analyse SNPs only in the core genome. Given much bacterial adaptability hinges on the accessory genome, this is an unsatisfactory limitation.

**Results:** We present a novel pan-genome graph structure and algorithms implemented in the software *pandora*, which approximates a sequenced genome as a recombinant of reference genomes, detects novel variation and then pan-genotypes multiple samples. The method takes fastq as input and outputs a multi-sample VCF with respect to an inferred data-dependent reference genome, and is available at https://github.com/rmcolq/pandora.

Constructing a reference graph from 578 *E. coli* genomes, we analyse a diverse set of 20 *E. coli* isolates. We show *pandora* recovers at least 13k more rare SNPs than single-reference based tools, achieves equal or better error rates with Nanopore as with Illumina data, 6-24x lower Nanopore error rates than other tools, and provides a stable framework for analysing diverse samples without reference bias. We also show that our inferred recombinant VCF reference genome is significantly better than simply picking the closest RefSeq reference.

**Conclusions:** This is a step towards comprehensive cohort analysis of bacterial pan-genomic variation, with potential impacts on genotype/phenotype and epidemiological studies.

## Background

Bacterial genomes evolve by multiple mechanisms including: mutation during replication, allelic and non-allelic homologous recombination. These processes result in a population of genomes that are mosaics of each other. Given multiple contemporary genomes, the segregating variation between them allows inferences to be made about their evolutionary history. These analyses are central to the study of bacterial genomics and evolution(1–4) with different questions requiring focus on separate aspects of the mosaic: fine-scale (mutations) or coarse (gene presence, synteny). In this paper, we provide a new and accessible conceptual model that combines both fine and coarse bacterial variation. Using this new understanding to better represent variation, we can access previously hidden single nucleotide polymorphisms (SNPs), insertions and deletions (indels).

Genes cover 85-90% of bacterial genomes(5), and shared gene content is commonly used as a measure of whole-genome similarity. In fact, the full set of genes present in a species - the *pan-genome* - is in general much larger than the number found in any single genome. A frequency distribution plot of genes within a set of bacterial genomes has a characteristic asymmetric U-shaped curve (6–10), as shown in Figure 1a. As a result, a collection of *Escherichia coli* genomes might only have 50% of their genes (and therefore their whole genome)(3) in common. This highlights a limitation in the standard approach to analysing genetic variation, whereby a single genome is treated as a reference, and all other genomes are interpreted as differences from it. In bacteria, a single reference genome will inevitably lack many of the genes in the pan-genome, and completely miss genetic variation therein (Figure 1b). We call this *hard reference bias*, to distinguish from the more common concern, that increased divergence of a reference from the genome under study leads to read-mapping problems, which we term *soft reference bias*. The standard workaround for these issues in bacterial genomics is to restrict analysis either to very similar genomes using a closely related reference (*e.g*. in an outbreak) or to analyse SNPs only in the core genome (present in most samples) and outside the core to simply study presence/absence of genes(11).

**Figure 1.**
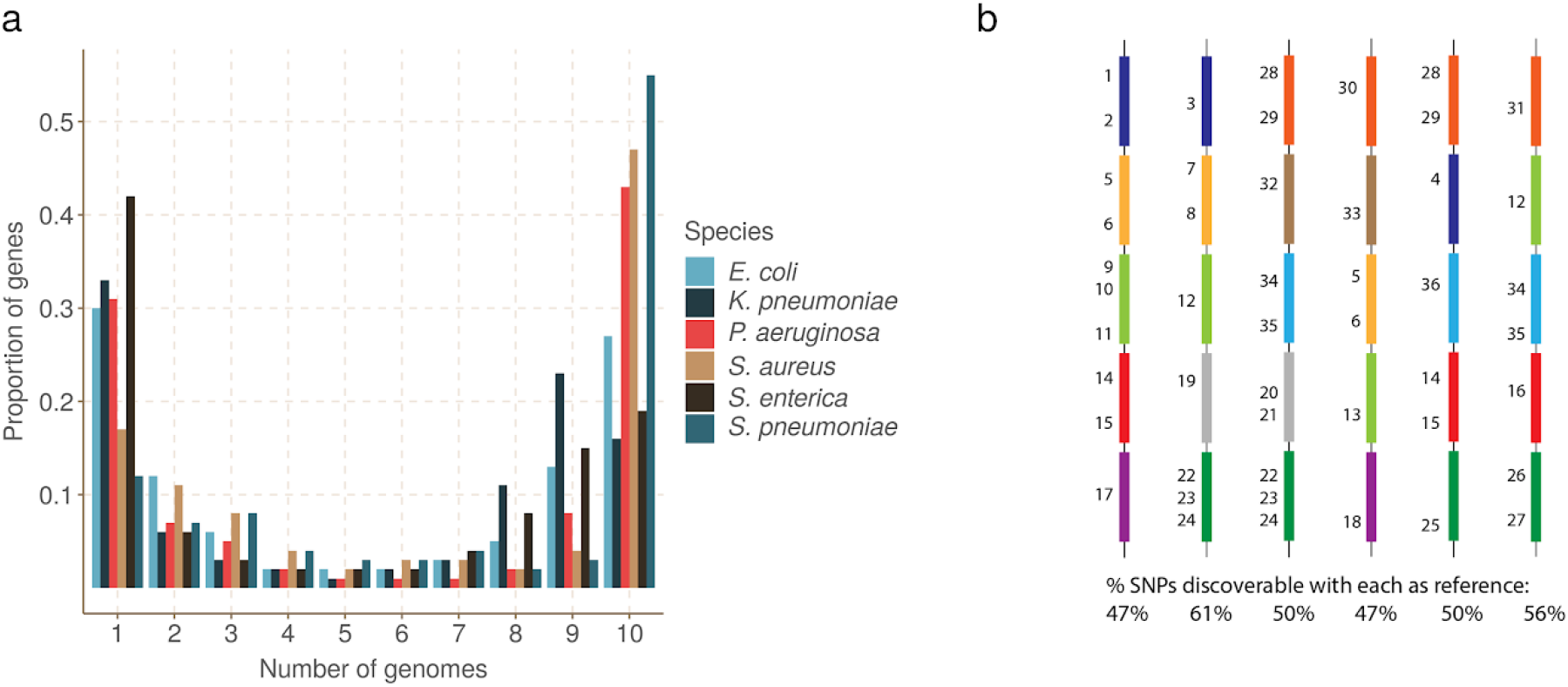
Universal gene frequency distribution in bacteria and the single-reference problem. a) Frequency distribution of genes in 10 genomes of 6 bacterial species (Escherischia coli, Klebsiella pneumoniae, Pseudomonas aeruginosa, Staphylococcus aureus, Salmonella enterica and Streptococcus pneumoniae) showing the characteristic U-shaped curve - most genes are rare or common. b) Illustrative depiction of the single-reference problem, a consequence of the U-shaped distribution. Each vertical column is a bacterial genome, and each coloured bar is a gene. Numbers are identifiers for SNPs - there are 50 in total. Thus the dark blue gene has 4 SNPs numbers 1-4. This figure does not detail which genome has which allele. Below each column is the proportion of SNPs that are discoverable when that genome is used as a reference genome. Because no single reference contains all the genes in the population, it can only access a fraction of the SNPs.

In this study we address the variation deficit caused by a single-reference approach. Given Illumina or Nanopore sequence data from potentially divergent isolates of a bacterial species, we attempt to detect all of the variants between them. Our approach is to decompose the pan-genome into atomic units (loci) which tend to be preserved over evolutionary timescales. Our loci are genes and intergenic regions in this study, but the method is agnostic to such classifications, and one could add any other grouping wanted (*e.g*. operons or mobile genetic elements). Instead of using a single genome as a reference, we collect a panel of representative reference genomes and use them to construct a set of reference graphs, one for each locus. Reads are mapped to this set of graphs and from this we are able to discover and genotype variation. By letting go of prior information on locus ordering in the reference panel, we are able to recognise and genotype variation in a locus regardless of its wider context. Since Nanopore reads are typically long enough to encompass multiple loci, it is possible to subsequently infer the order of loci - although that is outside the scope of this study.

The use of graphs as a generalisation of a linear reference is an active and maturing field(12–19). Much recent graph genome work has gone into showing that genome graphs reduce the impact of soft reference bias on mapping(12), and on generalising alignment to graphs(16,20). However there has not yet been any study (to our knowledge) addressing SNP analysis across a diverse cohort, including more variants that can fit on any single reference. In particular, all current graph methods require a reference genome to be provided in advance to output genetic variants in the standard Variant Call Format (VCF)(21) - thus immediately inheriting a hard bias when applied to bacteria (see Figure 1b).

We have made a number of technical innovations. First, a recursive clustering algorithm that converts a multiple sequence alignment (MSA) of a locus into a graph. This avoids the complexity “blowups” that plague graph genome construction from unphased VCF files(12,14). Second, a graph representation of genetic variation based on (w,k)-minimizers(22). Third, using this representation we avoid unnecessary full alignment to the graph and instead use quasi-mapping to genotype on the graph. Fourth, discovery of variation missing from the reference graph using local assembly. Fifth, use of a canonical dataset-dependent reference genome designed to maximise clarity of description of variants (the value of this will be made clear in the main text).

We describe these below, and evaluate our implementation, *pandora*, on a diverse set of *E. coli* genomes with both Illumina and Nanopore data. We show that, compared with reference-based approaches, *pandora* recovers a significant proportion of the missing variation in rare loci, performs much more stably across a diverse dataset, successfully infers a better reference genome for VCF output, and outperforms current tools for Nanopore data.

## Results

### Pan-genome graph representation

We set out to define a generalised reference structure which allows detection of SNPs and other variants across the whole pan-genome, without attempting to record long-range structure or coordinates. We define a *Pan-genome Reference Graph* (PanRG) as an unordered collection of sequence graphs, termed *local graphs*, each of which represents a locus, such as a gene or intergenic region. Each local graph is constructed from a MSA of known alleles of this locus, using a recursive cluster-and-collapse (RCC) algorithm (Supplementary Animation 1: recursive clustering construction). The output is guaranteed to be a directed acyclic sequence graph allowing hierarchical nesting of genetic variation while meeting a “balanced parentheses” criterion (see Figure 2b and Methods). Each path through the graph from source to sink represents a possible recombinant sequence for the locus. The disjoint nature of this pan-genome reference allows loci such as genes to be compared regardless of their wider genomic context. We implement this construction algorithm in the *make_prg* tool which outputs the graph as a file (see Figures 2a-c, Methods). Subsequent operations, based on this, are implemented in the software package *pandora*. The overall workflow is shown in Figure 2.

**Figure 2.**
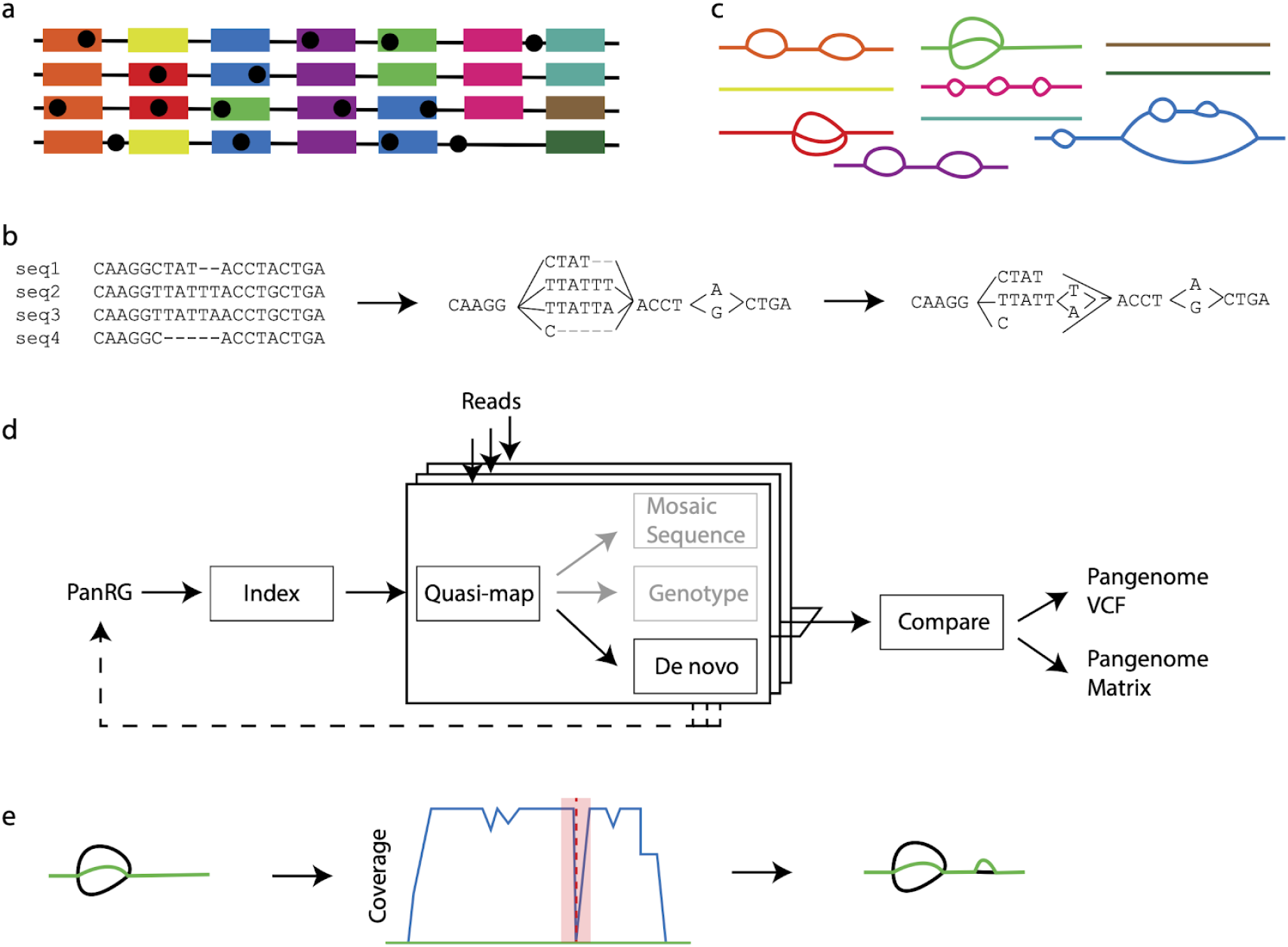
The *pandora* workflow. a) reference panel of genomes; colour signifies locus (gene or intergenic region) identifier, and blobs are SNPs. b) multiple sequence alignments (MSAs) for each locus are made and converted into a directed acyclic graph. c) local graphs constructed from the loci in the reference panel. d) Workflow: the collection of local graphs, termed the PanRG, is indexed. Reads from each sample under study are independently quasi-mapped to the graph, and a determination is made as to which loci are present in each sample. In this process, for each locus, a mosaic approximation of the sequence for that sample is inferred, and variants are genotyped. e) regions of low coverage are detected, and local de novo assembly is used to generate candidate novel alleles missing from the graph. Returning to d), the dotted line shows all the candidate alleles from all samples are then gathered and added to the MSAs at the start, and the PanRG is updated. Then, reads are quasi-mapped one more time, to the augmented PanRG, generating new mosaic approximations for all samples and storing coverages across the graphs; no de novo assembly is done this time. Finally, all samples are compared, and a VCF file is produced, with a per-locus reference that is inferred by pandora.

To index a PanRG, we generalise a type of sparse marker k-mer ((w,k)-minimizer), previously defined for strings, to directed acyclic graphs (see Methods). Each local graph is *sketched* with minimizing k-mers, and these are then used to construct a new graph (the k-mer graph) for each local graph from the PanRG. Each minimizing k-mer is a node, and edges are added between two nodes if they are adjacent minimizers on a path through the original local graph. This k-mer graph is isomorphic to the original if *w*≤*k* (and outside the first and last w+k-1 bases); all subsequent operations are performed on this graph, which, to avoid unnecessary new terminology, we also call the local graph.

A global index maps each minimizing k-mer to a list of all local graphs containing that k-mer and the positions therein. Long or short reads are approximately mapped (*quasi-mapped*) to the PanRG by determining the minimizing k-mers in each read. Any of these read quasi-mappings found in a local graph are called *hits*, and any local graph with sufficient clustered hits on a read is considered present in the sample.

### Initial sequence approximation as a mosaic of references

For each locus identified as present in a sample, we initially approximate the sample’s sequence as a path through the local graph. The result is a mosaic of sequences from the reference panel. This path is chosen to have maximal support by reads, using a dynamic programming algorithm on the graph induced by its (w,k)-minimizers (details in Methods). The result of this process serves as our initial approximation to the genome under analysis.

### Improved sequence approximation: modify mosaic by local assembly

At this point, we have quasi-mapped reads, and approximated the genome by finding the closest mosaic in the graph; however, we expect the genome under study to contain variants that are not present in the PanRG. Therefore, to allow discovery of novel SNPs and small indels that are not in the graph, for each sample and locus we identify regions of the inferred mosaic sequence where there is a drop in read coverage (as shown in Figure 2e). Slices of overlapping reads are extracted, and a form of *de novo* assembly is performed using a de Bruijn graph. Instead of trying to find a single correct path, the de Bruijn graph is traversed (see Methods for details) to all feasible candidate novel alleles for the sample. These alleles are added to the reference MSA for the locus, and the local graph is updated. If comparing multiple samples, the graphs are augmented with all new alleles from all samples at the same time.

### Optimal VCF-reference construction for multi-genome comparison

In the *compare* step of *pandora* (see Figure 2d), we enable continuity of downstream analysis by outputting genotype information in the conventional VCF(21). In this format, each row (record) describes possible alternative allele sequence(s) at a position in a (single) reference genome and information about the type of sequence variant. A column for each sample details the allele seen in that sample, often along with details about the support from the data for each allele.

To output graph variation, we first select a path through the graph to be the reference sequence and describe any variation within the graph with respect to this path as shown in Figure 3. We use the chromosome field to detail the local graph within the PanRG in which a variant lies, and the position field to give the position in the chosen reference path sequence for that graph. In addition, we output the reference path sequences used as a separate file.

**Figure 3.**
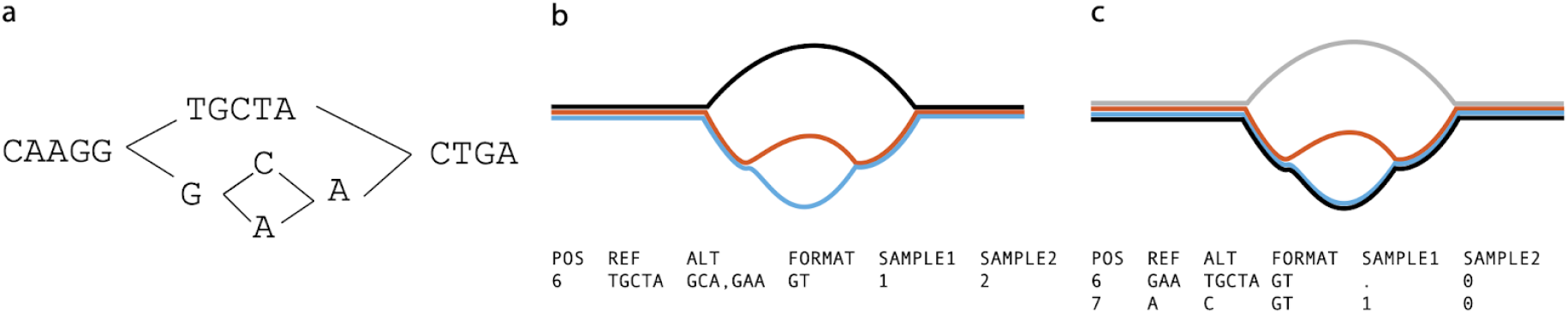
The representation problem. a) a local graph. b) The black allele is chosen as reference to enable representation in VCF. The blue/red SNP then requires flanking sequence in order to allow it to have a coordinate. The SNP is thus represented as two ALT alleles, each 3 bases long, and the user is forced to notice they only differ in one base. c) The blue path is chosen as the reference, thus enabling a more succinct and natural representation of the SNP.

For a collection of samples, we want small differences between samples to be recorded as short alleles in the VCF file rather than longer alleles with shared flanking sequence as shown in Figure 3b. We therefore choose the reference path for each local graph to be maximally close to the sample mosaic paths. To do this, we make a copy of the k-mer graph and increment the coverage along each sample mosaic path, producing a graph with higher weights on paths shared by more samples. We reuse the mosaic path-finding algorithm (see Methods) with a modified probability function defined such that the probability of a node is proportional to the number of samples covering it. This produces a dataset-dependent VCF reference able to succinctly describe segregating variation in the cohort of genomes under analysis.

### Constructing a PanRG of *E. coli*

We chose to evaluate *pandora* on the recombining bacterial species, *E. coli*, whose pan-genome has been heavily studied(7,23–26). MSAs for gene clusters curated with PanX(27) from 350 RefSeq assemblies were downloaded from http://pangenome.de on 3rd May 2018. MSAs for intergenic region clusters based on 228 *E. coli* ST131 genome sequences were previously generated with Piggy(28) for their publication. Whilst this panel of intergenic sequences does not reflect the full diversity within *E. coli*, we included them as an initial starting point. This resulted in an *E. coli* PanRG containing local graphs for 23,054 genes and 14,374 intergenic regions. *Pandora* took 24.4h in CPU time (2.3h in runtime with 16 threads) and 12.6 GB of RAM to index the PanRG. As one would expect from the U-shaped gene frequency distribution, many of the genes were rare in the 578 (=350+228) input genomes, and so 59%/44% of the genic/intergenic graphs were linear, with just a single allele.

### Constructing an evaluation set of diverse genomes

We first demonstrate that using a PanRG reduces hard bias when comparing a diverse set of 20 *E. coli* samples by comparison with standard single reference variant callers. We selected samples from across the phylogeny (including phylogroups A, B2, D and F(29)) where we were able to obtain both long and short read sequence data from the same isolate.

#### Two assemblies from phylogroup B1 are in the set of references, despite there being no sample in that phylogroup

We used Illumina-polished long read assemblies as truth data, masking positions where the Illumina data did not support the assembly (see Methods). As comparators, we used SAMtools(30) (the “classical” variant-caller based on pileups) and Freebayes(31) (a haplotype-based caller which reduces soft reference bias, wrapped by Snippy(32)) for Illumina data, and Medaka(33) and Nanopolish(34) for Nanopore data. In all cases, we ran the reference-based callers with 24 carefully selected reference genomes (see Methods, and Figure 4). We defined a “truth set” of 618,305 segregating variants by performing all pairwise whole genome alignments of the 20 truth assemblies, collecting SNP variants between the pairs, and deduplicating them by clustering into equivalence classes. Each class, or *pan-variant*, represents the same variant found at different coordinates in different genomes (see Methods). We evaluated error rate, pan-variant recall (PVR, proportion of truth set discovered) and average allelic recall (AvgAR, average of the proportion of alleles of each pan-variant that are found). To clarify the definitions, consider a toy example. Suppose we have three genes, each with one SNP between them. The first gene is rare, present in 2/20 genomes. The second gene is at an intermediate frequency, in 10/20 genomes. The third is a strict core gene, present in all genomes. The SNP in the first gene has alleles A,C at 50% frequency (1 A and 1 C). The SNP in the second gene has alleles G,T at 50% frequency (5 G and 5 T). The SNP in the third gene has alleles A,T with 15 A and 5 T. Suppose a variant caller found the SNP in the first gene, detecting the two correct alleles. For the second gene’s SNP, it detected only one G and one T, failing to detect either allele in the other 8 genomes. For the third gene’s SNP, it detected all the 5 T’s, but no A. Here, the pan-variant recall would be: (1 + 1 + 0) / 3 = 0.66 - *i.e*.score a 1 if both alleles are found, irrespective of how often- and the average allelic recall would be (2/2 + 2/10 + 5/20)/3=0.48.

**Figure 4.**
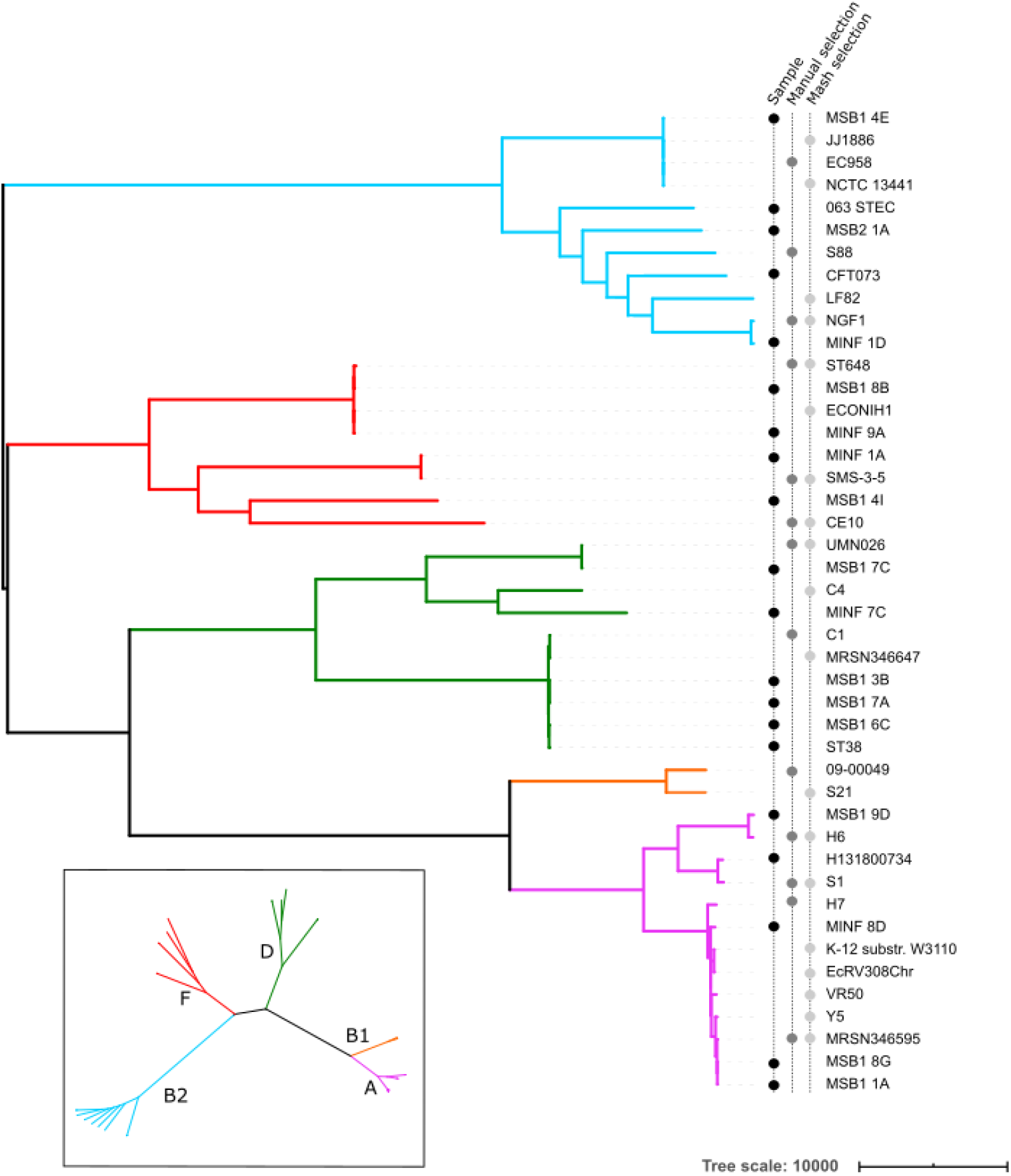
Phylogeny of 20 diverse E. coli along with references used for benchmarking single-reference variant callers. The 20 E. coli under study are labelled as samples in the left-hand of three vertical label-lines. Phylogroups (clades) are labelled by colour of branch, with the key in the inset. References were selected from RefSeq as being the closest to one of the 20 samples as measured by Mash, or manually selected from a tree (see Methods).

### Methylation-aware basecalling improves results

In Figure 5, we show for 4 samples the effect of methylation-aware Nanopore basecalling on the AvgAR/error rate curve for *pandora* with/without novel variant discovery via local assembly.

**Figure 5.**
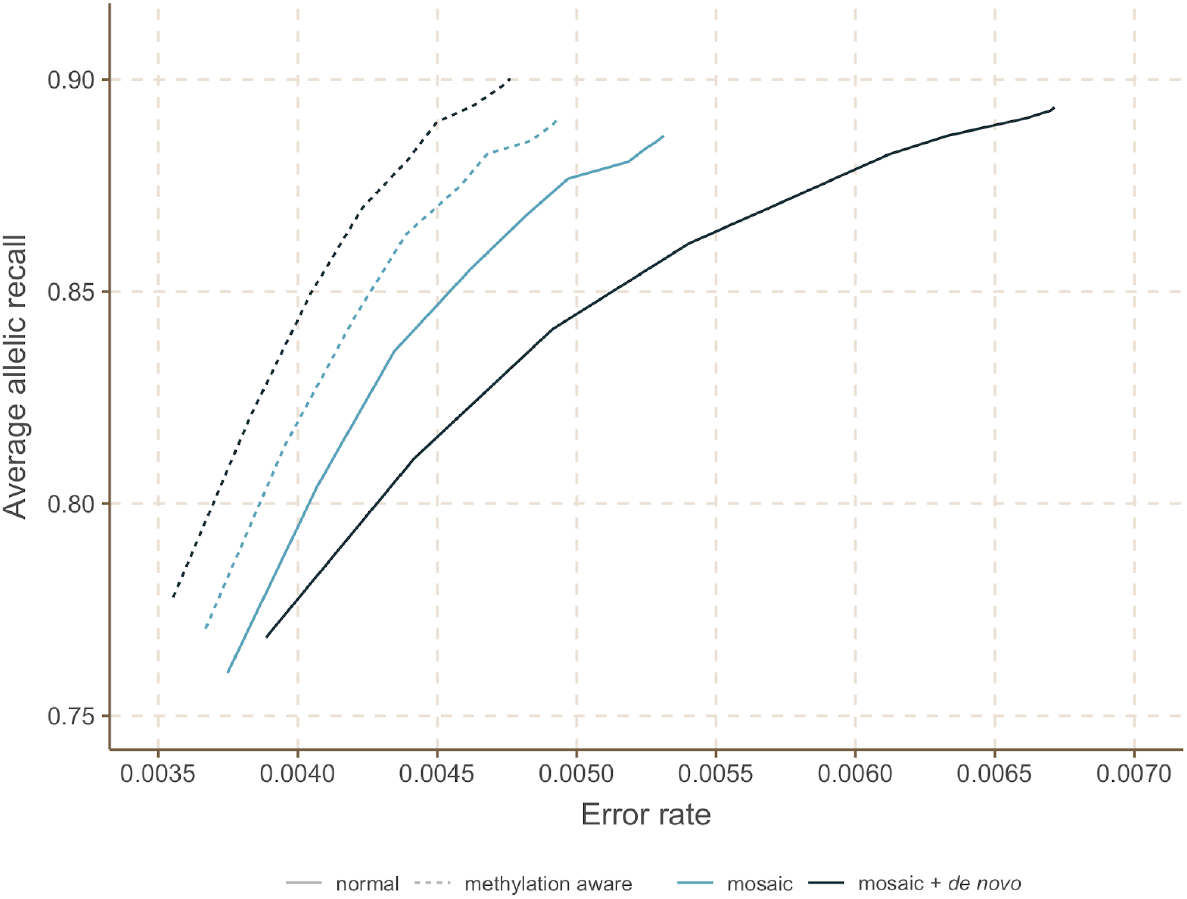
The effect of methylation-aware basecalling on local de novo assembly. We show the Average Allelic Recall and Error Rate curve for pandora with normal (solid line) or methylation-aware (dashed line) Guppy basecalling on 4 out of the 20 samples. For each of these input data, we show results for Pandora’s first approximation to a genome as a mosaic (recombinant) of the input reference panel (mosaic, light blue), and then the improved approximation with added de novo discovery (mosaic+de novo, dark blue).

The top right of each curve corresponds to completely unfiltered results; increasing the genotype confidence threshold (see Methods) moves each curve towards the bottom-left, increasing precision at the cost of recall. Notably, with normal basecalling, local *de novo* assembly increases the error rate from 0.53% to 0.67%, with a negligible increase in recall, from 88.7% to 89.3%, whereas with methylation-aware basecalling it increases the recall from 89.1% to 90% and slightly decreases the error rate from 0.49% to 0.48%. On the basis of this, from here on we work entirely with reads that are basecalled with a methylation-aware model, and move to the full dataset of 20 samples.

### Benchmarking recall, error rate and dependence on reference

We show in Figures 6a,b the Illumina and Nanopore AvgAR/recall plots for *pandora* and four single-reference tools with no filters applied. For all of these, we modify only the minimum genotype confidence to move up and down the curves (see Methods).

**Figure 6.**
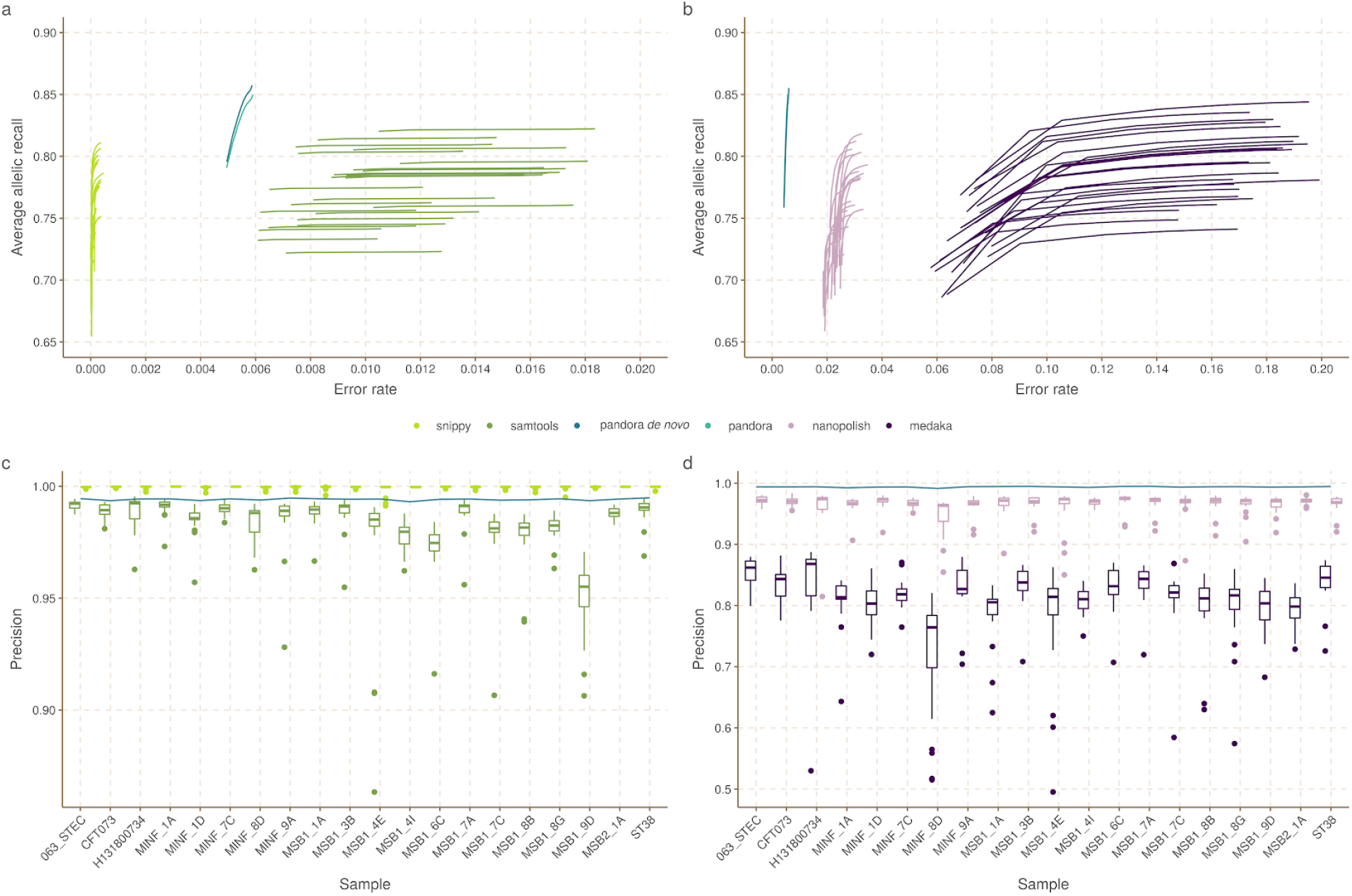
Benchmarks of recall/error and dependence of precision on reference genome, for *pandora* and other tools on 20-way dataset. a) The average allelic recall and error rate curve for pandora, SAMtools and snippy on 100x of Illumina data. Snippy/SAMtools both run 24 times with the different reference genomes shown in figure 4, resulting in multiple lines for each tool (one for each reference) b) The average allelic recall and error rate curve for pandora, medaka and nanopolish on 100x of Nanopore data; multiple lines for medaka/nanopolish, one for each reference genome. Note panels a and b have the same y axis scale and limits, but different × axes; c) The precision of pandora, SAMtools and snippy on 100x of Illumina data. The boxplots show the distribution of SAMtools’ and snippy’s precision depending on which of the 24 references was used, and the blue line connects pandora’s results; d) The precision of pandora (line plot), medaka and nanopolish (both boxplots) on 100x of Nanopore data. Note different y axis scale/limits in panels c,d.

We highlight three observations. Firstly, *pandora* achieves essentially the same recall and error rate for the Illumina and Nanopore data (85% AvgAR and 0.6% error rate at the top-right of the curve, completely unfiltered). Second, choice of reference has a significant effect on both AvgAR and error rate for the single-reference callers; the reference which enables the highest recall does not lead to the best error rate (for *SAMtools* and *medaka* in particular). Third, *pandora* achieves better AvgAR (86%) than all other tools (all between 81% and 84%, see Supplementary Table 2), and a better error rate (0.6%) than *SAMtools* (1%), *nanopolish* (2.4%) and *medaka* (14.8%). However, *snippy* achieves a significantly better error rate than all other tools (0.01%). We confirmed that adding further filters slightly improved error rates, but did not change the overall picture (Supplementary Figure 1, Methods, Supplementary Table 2). The results are also in broad agreement if the PVR is plotted instead of AvgAR (Supplementary Figure 2). However, these AvgAR and PVR figures are hard to interpret because *pandora* and the reference-based tools have recall that varies differently across the locus frequency spectrum - we explore this further below.

We ascribe the similarity between the Nanopore and Illumina performance of *pandora* to three reasons. First, the PanRG is a strong prior - our first approximation does not contain any Nanopore sequence, but simply uses quasi-mapped reads to find the nearest mosaic in the graph. Second, mapping long Nanopore reads which completely cover entire genes is easier than mapping Illumina data, and allows us to filter out erroneous k-mers within reads after deciding when a gene is present. Third, this performance is only achieved when we use methylation-aware basecalling of Nanopore reads, presumably removing most systematic bias (see Figure 5).

In Figure 6c,d we show for Illumina and Nanopore data, the impact of reference choice on the precision of calls on each of the 20 samples. While precision is consistent across all samples for *pandora*, we see a dramatic effect of reference-choice on precision of *SAMtools, medaka* and *nanopolish*. The effect is also detectable for *snippy*, but to a much lesser extent.

Finally, we measured the performance of locus presence detection, restricting to genes/intergenic regions in the PanRG, so that in principle perfect recall would be possible (see Methods). In Supplementary Figure 3 we show the distribution of locus presence calls by *pandora*, split by length of locus for Illumina and Nanopore data. Overall, 93.8%/94.3% of loci were correctly classified as present or absent for Illumina/Nanopore respectively. Misclassifications were concentrated on small loci (below 500bp). While 59.2%/57.4% of all loci in the PanRG are small, 75.5%/74.8% of false positive calls and 98.7%/98.1% of false negative calls are small loci (see Supplementary Figure 3).

### *Pandora* detects rare variation inaccessible to single-reference methods

Next, we evaluate the key deliverable of *pandora* - the ability to access genetic variation within the accessory genome. We plot this in Figure 7, showing PVR of SNPs in the truth set which overlap genes or intergenic regions from the PanRG, broken down by the number of samples the locus is present in.

**Figure 7.**
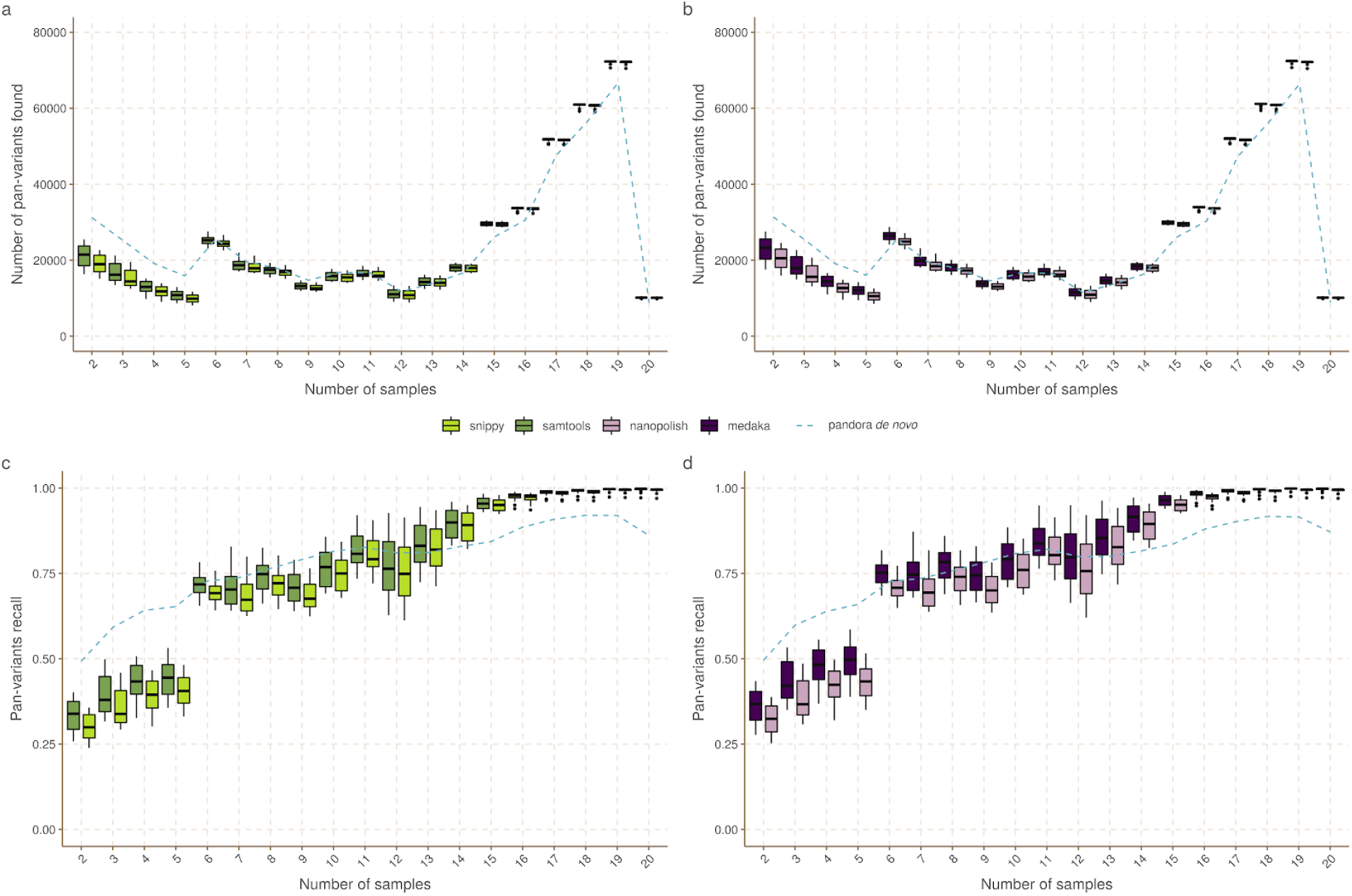
Pan-variant recall across the locus frequency spectrum. Every SNP occurs in a locus, which is present in some subset of the full set of 20 genomes. In all panels the SNPs in the golden truth set are broken down by the number of samples the locus is present in. Left panels (a, c) show results for pandora (dotted line), snippy and SAMtools with Illumina data. Right panels (b, d) show results for pandora, nanopolish and medaka with Nanopore data. Top panels (a, b) show the absolute count of pan-variants found; Bottom panels (c, d) show the proportion of pan-variants found

If we restrict our attention to rare variants (present only in 2-5 genomes), we find *pandora* recovers at least 19644/26674/13108/22331 more SNPs than *SAMtools/snippy/medaka/nanopolish* respectively. As a proportion of rare SNPs in the truth set, this is a lift in PVR of 12/17/8/14% respectively. If, instead of pan-variant recall, we look at the variation of AvgAR across the locus frequency spectrum (see Supplementary Figure 4), the gap between *pandora* and the other tools on rare loci is even larger. These observations, and Figure 6, confirm and quantify the extent to which we are able to recover accessory genetic variation that is inaccessible to single-reference based methods.

### *Pandora* has consistent results across *E. coli* phylogroups

We measure the impact of reference bias (and population structure) by quantifying how recall varies in phylogroups A, B2, D, and F dependent on whether the reference genome comes from the same phylogroup.

We plot the results for *snippy* with 5 exemplar references in Figure 8a (results for all tools and for all references are in Supplementary Figures 5-8), showing that single references give 5-10% higher recall for samples in their own phylogroup than other phylogroups. By comparison, *pandora*’s recall is much more consistent, staying stable at ∼89% for all samples regardless of phylogroup. References in phylogroups A and B2 achieve higher recall in their own phylogroup, but consistently worse than *pandora* for samples in the other phylogroups (in which the reference does not lie). References in the external phylogroup B1, for which we had no samples in our dataset, achieve higher recall for samples in the nearby phylogroup A (see inset, Figure 4), but lower than *pandora* for all others. We also see that choosing a reference genome from phylogroup F (red), which sits intermediate to the other phylogroups, provides the most uniform recall across other groups - 2-5% higher than *pandora*.

**Figure 8.**
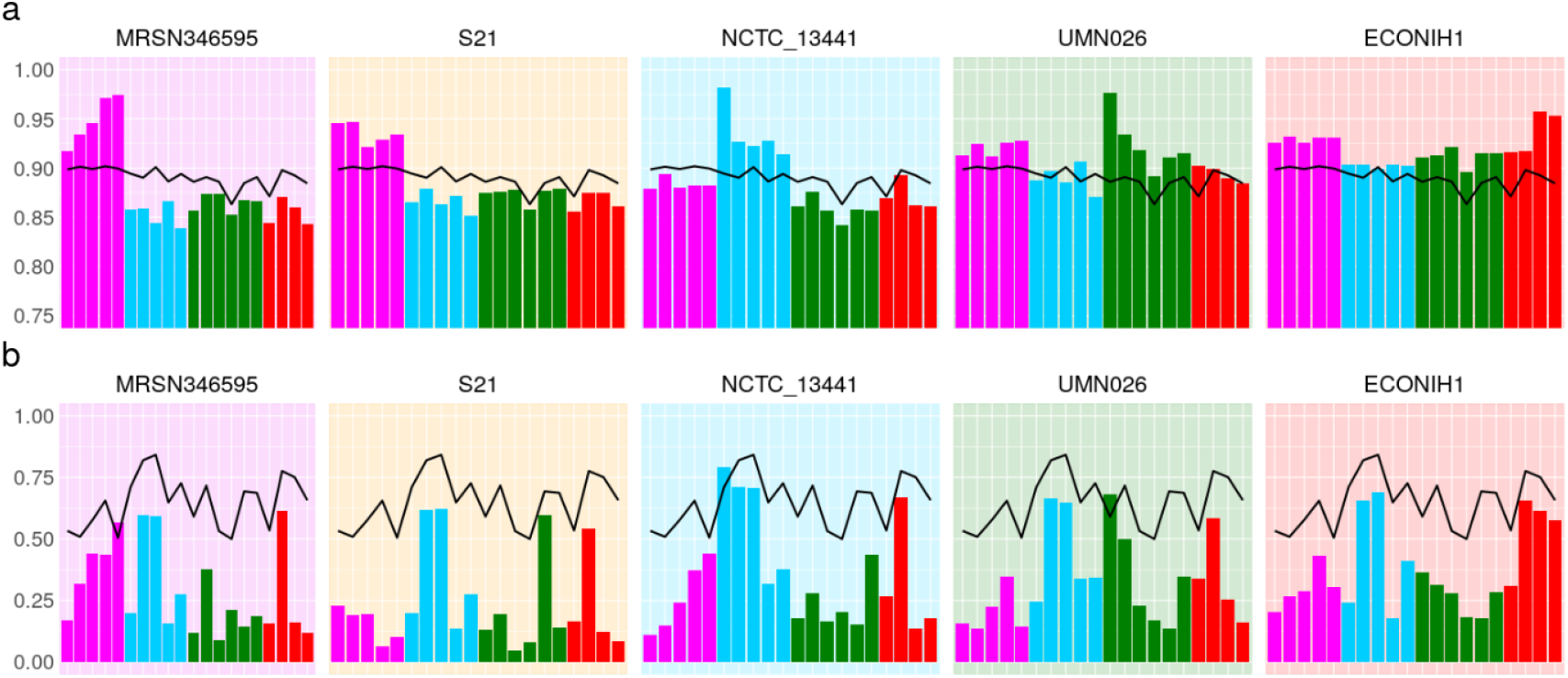
Single reference callers achieve higher recall for samples in the same phylogroup as the reference genome, but not for rare loci. a) pandora recall (black line) and snippy recall (coloured bars) on the 20 samples; each histogram corresponds to the use of one of 5 exemplar references, one from each phylogroup. The background colour denotes the reference’s phylogroup (see Figure 4 inset); note that phylogroup B1 (yellow background) is an outgroup, containing no samples in this dataset; b) Same as a) but restricted to SNPs present in precisely two samples (i.e. where 18 samples have neither allele because the entire locus is missing). Note the differing y-axis limits in the two panels.

These results will, however, be dominated by the shared, core genome. If we replot Figure 8a, restricting to variants in loci present in precisely 2 genomes (abbreviated to 2-variants; Figure 8b), we find that *pandora* achieves 50-84% recall for each sample (complete data in Supplementary Figure 9). By contrast, for any choice of reference genome, the results for single-reference callers vary dramatically per sample. Most samples have recall under 25%, and there is no pattern of improved recall for samples in the same phylogroup as the reference. Following up that last observation, if we look at which pairs of genomes share 2-variants (Figure 9), we find there is no enrichment within phylogroups at all. This simply confirms in our data that presence of rare loci is not correlated with the overall phylogeny.

**Figure 9.**
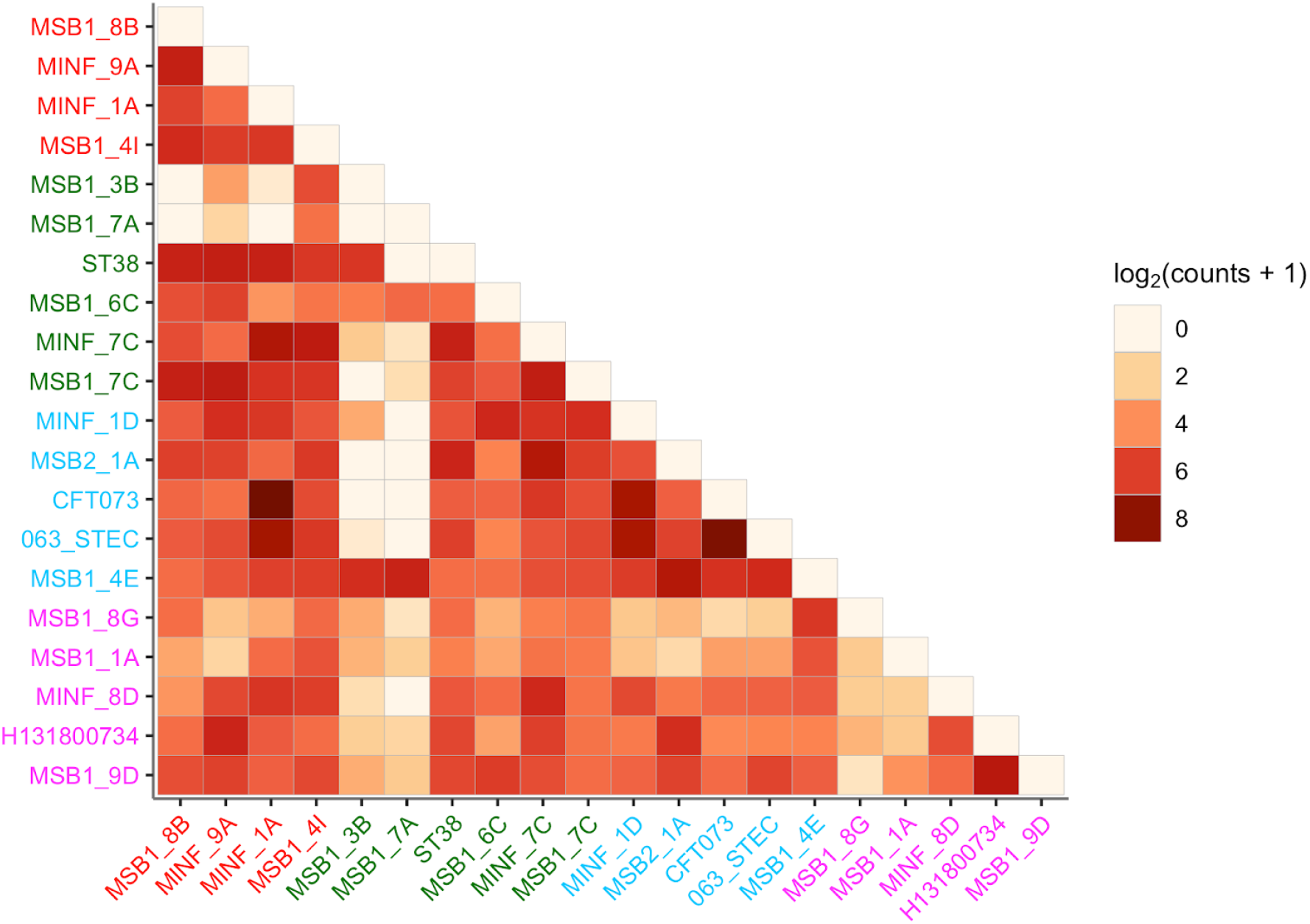
Sharing of variants present in precisely 2 genomes, showing which pairs of genomes they lie in and which phylogroups; darker colours signify higher counts (log scale). Genomes are coloured by their phylogroup (see Figure 4 inset).

### *Pandora* VCF reference is closer to samples than any single reference

The relationship between phylogenetic distance and gene repertoire similarity is not linear. In fact, 2 genomes in different phylogroups may have more similar accessory genes than 2 in the same phylogroup - as illustrated in the previous section (also see Figure 3 in Rocha(3)). As a result, it is unclear *a priori* how to choose a good reference genome for comparison of accessory loci between samples. *Pandora* specifically aims to construct an appropriate reference for maximum clarity in VCF representation. We evaluate how well *pandora* is able to find a VCF reference close to the samples under study as follows. We first identified the location of all loci in all the 20 sample assemblies and the 24 references (see Methods).

We then measured the edit distance between each locus in each of the references and the corresponding version in the 20 samples. We found that the *pandora*’s VCF-reference lies within 1% edit distance (scaled by locus length) of the sample far more than any of the references for loci present in <=14 samples (Figure 10; note the log scale). The improvement is much reduced in the core genome; essentially, in the core, a phylogenetically close reference provides a good approximation, but it is hard to choose a single reference that provides a close approximation to all rare loci. By contrast, *pandora* is able to leverage its reference panel, and the dataset under study, to find a good approximation.

**Figure 10.**
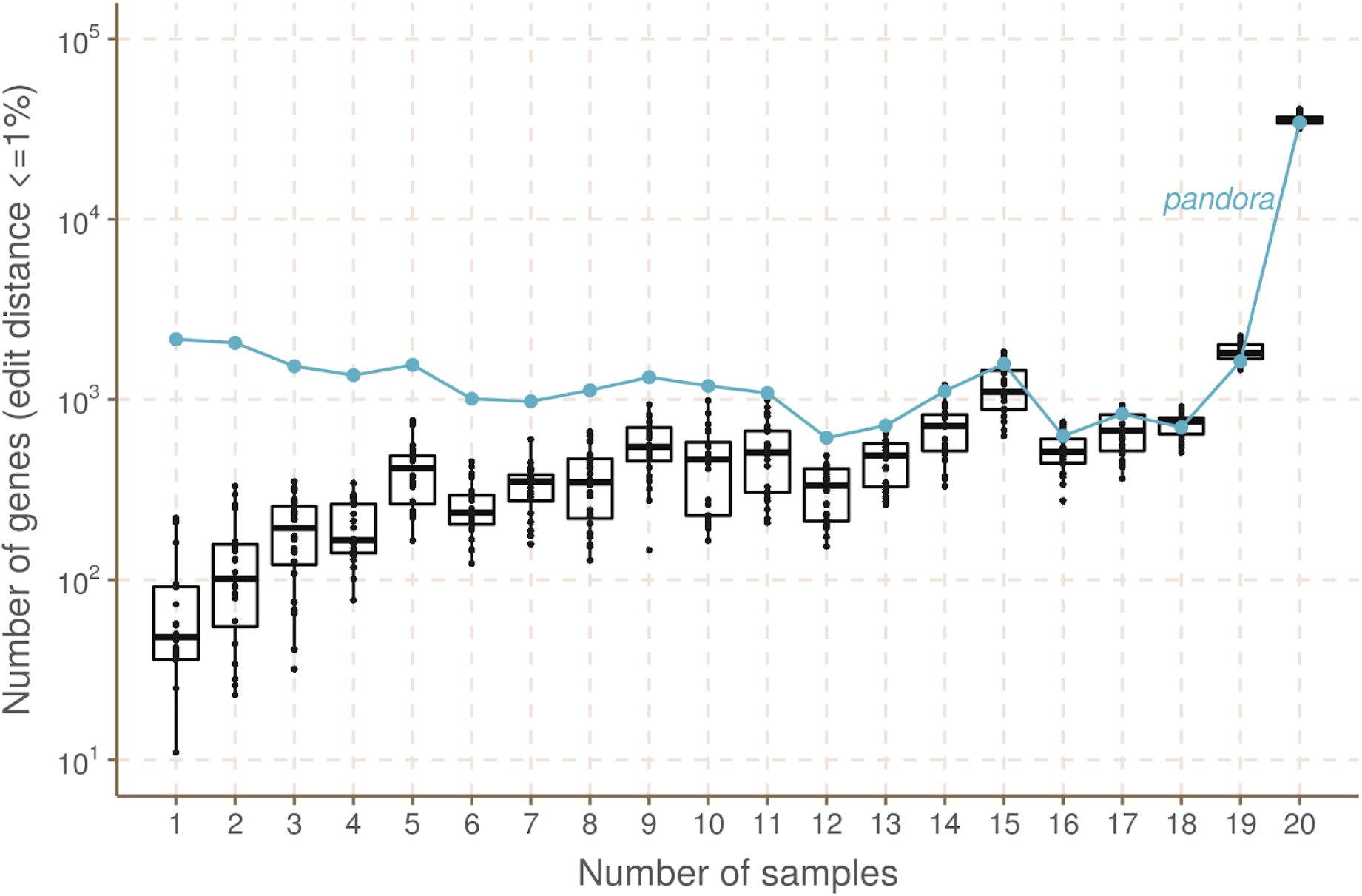
How often do references closely approximate a sample? pandora aims to infer a reference for use in its VCF, which is as close as possible to all samples. We evaluate the success of this here. The x-axis shows the number of genomes in which a locus occurs. The y-axis shows the (log-scaled) count of loci in the 20 samples that are within 1% edit distance (scaled by locus length) of each reference - box plots for the reference genomes, and line plot for the VCF reference inferred by pandora.

### Computational performance

Performance measurements for single-sample analysis by *pandora* and benchmarked tools are shown in Supplementary Table 3. In short, *pandora* took 3-4 hours per sample (using 16 cores and up to 10.7 GB of RAM), which was slower than *snippy* (0.1h, 4 cores), *SAMtools* (0.3h, 1 core) and *medaka* (0.3h, 4 cores), but faster than *nanopolish* (4.6h, 16 cores).

*Pandora* alone can do joint analysis of multiple samples and this is currently the most expensive *pandora* step. Parallelising by gene on a compute cluster, it took 8 hours to augment the PanRG with novel alleles. This was dominated by the Python implementation of the RCC clustering algorithm (see Methods) and the use of Clustal Omega(35) for MSA. 90% of loci required less than 30 minutes to process, and the remainder took less than 2 hours (see Methods). We discuss below how this could be improved. Finally, it took 28/46 hours to compare the samples (produce the joint VCF file) for Illumina/Nanopore. Mapping comprised ∼10% of the Illumina time, and ∼50% of the Nanopore time. Dynamic programming and genotyping the VCF file took ∼90% of the Illumina time, and ∼50% of the Nanopore time.

## Discussion

Bacteria are the most diverse and abundant cellular life form(36). Some species are exquisitely tuned to a particular niche (e.g. obligate pathogens of a single host) while others are able to live in a wide range of environments (e.g. *E. coli* can live on plants, in the earth, or commensally in the gut of various hosts). Broadly speaking, a wider range of environments correlates with a larger pan-genome, and some parts of the gene repertoire are associated with specific niches(37). Our perception of a pan-genome therefore depends on our sampling of the unknown underlying population structure, and similarly the effectiveness of a PanRG will depend on the choice of reference panel from which it is built.

Many examples from different species have shown that bacteria are able to leverage this genomic flexibility, adapting to circumstance sometimes by using or losing novel genes acquired horizontally, and at other times by mutation. There are many situations where precise nucleotide-level variants matter in interpreting pan-genomes. Some examples include: compensatory mutations in the chromosome reducing the fitness burden of new plasmids(38–40); lineage-specific accessory genes with SNP mutations which distinguish carriage from infection(41); SNPs within accessory drug resistance genes leading to significant differences in antibiograms(42); and changes in CRISPR spacer arrays showing immediate response to infection(43,44). However, up until now there has been no automated way of studying non-core gene SNPs at all; still less a way of integrating them with gene presence/absence information. *Pandora* solves these problems, allowing detection and genotyping of core and accessory variants. It also addresses the problem of what reference to use as a coordinate system, inferring a mosaic “VCF reference” which is as close as possible to all samples under study. We find this gives more consistent SNP-calling than any single reference in our diverse dataset. We focussed primarily on Nanopore data when designing *pandora*, and show it is possible to achieve higher quality SNP calling with this data than with current Nanopore tools. Together, these results open the door for empirical studies of the accessory genome, and for new population genetic models of the pan-genome from the perspective of both SNPs and gene gain/loss.

Prior graph genome work, focussing on soft reference bias (in humans), has evaluated different approaches for selecting alleles for addition to a population graph, based on frequency, avoiding creating new repeats, and avoiding exponential blowup of haplotypes in clusters of variants(45). This approach makes sense when you have unphased diploid VCF files and are considering all recombinants of clustered SNPs as possible. However, this is effectively saying we consider the recombination rate to be high enough that all recombinants are possible. Our approach, building from local MSAs and only collapsing haplotypes when they agree for a fixed number of bases, preserves more haplotype structure and avoids combinatorial explosion. Another alternative approach was recently taken by Norri *et al*.(46), inferring a set of pseudo founder genomes from which to build the graph.

Another issue is how to select the reference panel of genomes in order to minimize hard reference bias. One cannot escape the U-shaped frequency distribution; whatever reference panel is chosen, future genomes under study will contain rare genes not present in the PanRG. Given the known strong population structure in bacteria, and the association of accessory repertoires with lifestyle and environment, we would advocate sampling by geography, host species (if appropriate), lifestyle (e.g. pathogenic versus commensal) and/or environment. In this study we built our PanRG from a biassed dataset (RefSeq) which does not attempt to achieve balance across phylogeny or ecology, limiting our pan-variant recall to 49% for rare variants (see Figure 7c,d). A larger, carefully curated input panel, such as that from Horesh et al(47), would provide a better foundation and potentially improve results.

A natural question is then to ask if the PanRG should continually grow, absorbing all variants ever encountered. From our perspective, the answer is no - a PanRG with variants at all non-lethal positions would be potentially intractable. The goal is not to have every possible allele in the PanRG - no more than a dictionary is required to contain absolutely every word that has ever been said in a language. As with dictionaries, there is a trade-off between completeness and utility, and in the case of bacteria, the language is far richer than English. The perfect PanRG contains the vast majority of the genes and intergenic regions you are likely to meet, and just enough breadth of allelic diversity to ensure reads map without too many mismatches. Missing alleles should be discoverable by local assembly and added to the graph, allowing multi-sample comparison of the cohort under study. This allows one to keep the main PanRG lightweight enough for rapid and easy use.

We finish with three potential applications of *pandora*. First, the PanRG should provide a more interpretable substrate for pan-genome-wide Genome-Wide Association Studies, as current methods are forced to either ignore the accessory genome or reduce it to k-mers or unitigs(48–50). Second, if performing prospective surveillance of microbial isolates taken in a hospital, the PanRG provides a consistent and unchanging reference, which will cope with the diversity of strains seen without requiring the user to keep switching reference genome. In a sense it behaves similarly to whole-genome Multi-Locus Sequence Typing (wgMLST)(51), with more flexibility, support for intergenic regions, and without the all-or-nothing behaviour when alleles have a novel SNP. Third, if studying a fixed dataset very carefully, then one may not want to use a population PanRG, as it necessarily will miss some rare accessory genes in the dataset. In these circumstances, one could construct a reference graph purely of the genes/intergenic regions present in this dataset.

There are a number of limitations to this study. Firstly, *pandora* is not yet a fully-fledged production tool. There are two steps that constitute bottlenecks in terms of RAM and speed. The RCC algorithm used for local graph construction is currently implemented in Python. However, the underlying algorithm is amenable to a much higher performance implementation, which is now in progress. Also, we use Clustal Omega(35) for the MSA stage, and there are faster options which we could use, including options for augmenting an MSA without a complete rebuild (e.g. MAFFT), which is exactly what we need after local assembly discovers novel alleles. Secondly, we do not see any fundamental reason why the *pandora* error rate should be worse than Snippy on Illumina data (see Figure 6C), and will be working to improve this. Finally, by working in terms of atomic loci instead of a monolithic genome-wide graph, *pandora* opens up graph-based approaches to structurally diverse species (and eases parallelisation) but at the cost of losing genome-wide ordering. At present, ordering can be resolved by (manually) mapping *pandora*-discovered genes onto whole genome assemblies. However the design of *pandora* also allows for gene-ordering inference: when Nanopore reads cover multiple genes, the linkage between them is stored in a secondary de Bruijn graph where the alphabet consists of gene identifiers. This results in a huge alphabet, but the k-mers are almost always unique, dramatically simplifying “assembly” compared with normal DNA de Bruijn graphs. This work is still in progress and the subject of a future study. In the meantime, *pandora* provides new ways to access previously hidden variation.

## Conclusions

The algorithms implemented in *pandora* provide, to our knowledge, the first solution to the problem of analysing core and accessory genetic variation across a set of bacterial genomes. This study demonstrates as good SNP genotype error rates with Nanopore as with Illumina data and improved recall of accessory variants. It also shows the benefit of an inferred VCF reference genome over simply picking from RefSeq. The main limitations were the use of a biassed reference panel (RefSeq) for building the PanRG, and the comparatively slow performance of one module, currently implemented in Python - both of which are addressable, not fundamental limitations. This opens the door to improved analyses of many existing and future bacterial genomic datasets.

## Methods

### Local graph construction

We construct each local graph in the PanRG from an MSA using an iterative partitioning process. The resulting sequence graph contains nested bubbles representing alternative alleles.

Let *A* be an MSA of length *n*. For each row of the MSA *a* = {*a*_0_,…, *a*_*n* −1_} ∈ *A* let *a*_*i,j*_ = {*a*_*i*_,…, *a*_*j*−1_} be the subsequence of *a* in interval [*i, j*). Let *s*(*a*) be the DNA sequence obtained by removing all non-AGCT symbols. We can partition alignment *A* either *vertically* by partitioning the interval [0, *n*) or *horizontally* by partitioning the set of rows of *A*. In both cases, the partition induces a number of sub-alignments.

For vertical partitions, we define *slice*_*A*_(*i, j*) = {*a*_*i,j*_: *a* ∈ *A*}. We say that interval [*i, j*) is a *match* interval if *j i* ≥ *m*, where *m* = 7 is the default minimum match length, and there is a single non-trivial sequence in the slice, i.e. |{*s*(*a*): ∈ *slice* _*A*_ (*i, j*) *and s*(*a*) ≠ “”}|=1. Otherwise, we call it a *non-match* interval.

For horizontal partitions, we use *K* -means clustering(52) to divide sequences into increasing numbers of clusters *K* = 2, 3,… until the *inertia*, a measure of the within-cluster diversity, is half that of the original full set of sequences. More formally, let *U* be the set of all *m*-mers (substrings of length *m*, the minimum match length) in {*s*(*a*): *a* ∈ *A*}. For *a* ∈ *A* we transform sequence *s*(*a*) into a count vector 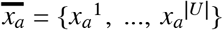 where *x*_*a*_^*i*^ are the counts of the unique *m*-mers in *U*. For *K* clusters 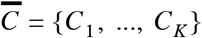, the inertia is defined as

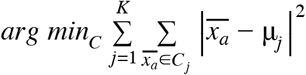

where 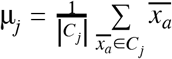 is the mean of cluster *j*.

The recursive algorithm first partitions an MSA vertically into match and non-match intervals. Match intervals are *collapsed* down to the single sequence they represent. Independently for each non-match interval, the alignment slice is partitioned horizontally into clusters. The same process is then applied to each induced sub-alignment until a maximum number of recursion levels, *r* = 5, has been reached. For any remaining alignments, a node is added to the local graph for each unique sequence. See Supplementary Animation 1 to see an example of this algorithm. We name this algorithm Recursive Cluster and Collapse (RCC), and implement in the make_prg repository (see Code Availability).

### (w,k)-minimizers of graphs

We define (w,k)-minimizers of strings as in Li (2016) (53). Let φ : Σ^*k*^ → ℜ be a k-mer hash function and let π : Σ^*^ × {0, 1} → Σ^*^ be defined such that π(*s*, 0) = *s* and 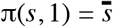, where 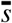 is the reverse complement of *s*. Consider any integers *k* ≥ *w* > 0. For window start position 0 ≤ *j* ≤ |*s*| *w k* + 1, let

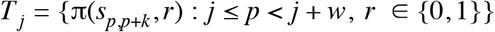

be the set of forward and reverse-complement k-mers of *s* in this window. We define a (w,k)-minimizer to be any triple (*h, p, r*) such that

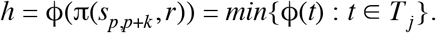

The set *W* (*s*) of (w,k)-minimizers for *s*, is the union of minimizers over such windows.

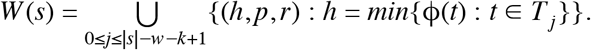

We extend this definition intuitively to an acyclic sequence graph G = (V,E). Define |*v*| to be the length of the sequence associated with node *v* ∈ *V* and let *i* = (*v, a, b*), 0 ≤ *a* ≤ *b* ≤ |*v*| represent the sequence interval [a,b) on v. We define a *path* in G by

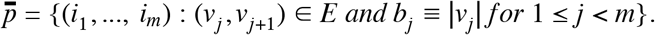

This matches the intuitive definition for a path in a sequence graph except that we allow the path to overlap only part of the sequence associated with the first and last nodes. We will use 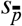 to refer to the sequence along the path 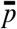 in the graph.

Let 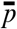 be a path of length w+k-1 in G. The string 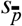 contains w consecutive k-mers for which we can find the (w,k)-minimizer(s) as before. We therefore define the (w,k)-minimizer(s) of the graph G to be the union of minimizers over all paths of length w+k-1 in G:

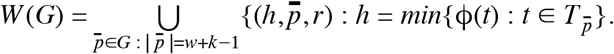

### Local graph indexing with (w,k)-minimizers

To find minimizers for a graph we use a streaming algorithm as described in Supplementary Algorithm 1. For each minimizer found, it simply finds the next minimizer(s) until the end of the graph has been reached.

Let *walk*(*v, i, w, k*) be a function which returns all vectors of w consecutive k-mers in G starting at position i on node v. Suppose we have a vector of k-mers x. Let *shift*(*x*) be the function which returns all possible vectors of k-mers which extend × by one k-mer. It does this by considering possible ways to walk one letter in G from the end of the final k-mer of x. For a vector of k-mers of length w, the function *minimize*(*x*) returns the minimizing k-mers of x.

We define K to be a *k-mer graph* with nodes corresponding to minimizers 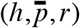. We add edge (u,v) to K if there exists a path in G for which u and v are both minimizers and v is the first minimizer after u along the path. Let *K* ← *add*(*s, t*) denote the addition of nodes s and t to K and the directed edge (s,t). Let *K* ← *add*(*s, T*) denote the addition of nodes *s* and *t* ∈ *T* to K as well as directed edges (s,t) for *t* ∈ *T*, and define *K* ← *add*(*S, t*) similarly.

The resulting PanRG index stores a map from each minimizing k-mer hash value to the positions in all local graphs where that (w,k)-minimizer occurred. In addition, we store the induced k-mer graph for each local graph.

### Quasi-mapping reads

We infer the presence of PanRG loci in reads by quasi-mapping. For each read, a sketch of (w,k)-minimizers is made, and these are queried in the index. For every (w,k)-minimizer shared between the read and a local graph in the PanRG index, we define a *hit* to be the coordinates of the minimizer in the read and local graph and whether it was found in the same or reverse orientation. We define clusters of hits from the same read, local graph and orientation if consecutive read coordinates are within a certain distance. If this cluster is of sufficient size, the locus is deemed to be present and we keep the hits for further analysis. Otherwise, they are discarded as noise. The default for this “sufficient size” is at least 10 hits and at least 1/5th the length of the shortest path through the k-mer graph (Nanopore) or the number of k-mers in a read sketch (Illumina). Note that there is no requirement for all these hits to lie on a single path through the local graph. A further filtering step is therefore applied after the sequence at a locus is inferred to remove false positive loci, as indicated by low mean or median coverage along the inferred sequence by comparison with the global average coverage. This quasi-mapping procedure is described in pseudocode in Supplementary Algorithm 2.

### Initial sequence approximation as a mosaic of references

For each locus identified as present in the set of reads, quasi-mapping provides (filtered) coverage information for nodes of the directed acyclic k-mer graph. We use these to approximate the sequence as a mosaic of references as follows. We model k-mer coverage with a negative binomial distribution and use the simplifying assumption that k-mers are read independently. Let Θ be the set of possible paths through the k-mer graph, which could correspond to the true genomic sequence from which reads were generated. Let r + s be the number of times the underlying DNA was read by the machine, generating a k-mer coverage of s, and r instances where the k-mer was sequenced with errors. Let 1 − p be the probability that a given k-mer was sequenced correctly. For any path θ ∈ Θ, let {*X*_1_,…, *X*_*M*_} be independent and identically distributed random variables with probability distribution 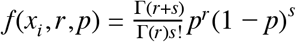, representing the k-mer coverages along this path. Since the mean and variance are 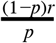 and 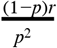 we solve for r and p using the observed k-mer coverage mean and variance across all k-mers in all graphs for the sample. Let *D* be the k-mer coverage data seen in the read dataset. We maximise the log-likelihood-inspired score 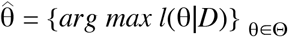 where 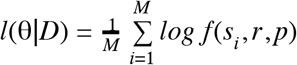 is the observed coverage of the *i* -th k-mer in θ. By construction, the k-mer graph is directed and acyclic so this maximisation problem can be solved with a dynamic programming algorithm (for pseudocode, see Supplementary Algorithm 3).

For choices of w ≤ k there is a unique sequence along the discovered path through the k-mer graph (except in rare cases within the first or last w-1 bases). We use this closest mosaic of reference sequences as an initial approximation of the sample sequence.

### *De novo* variant discovery

The first step in our implementation of local *de novo* variant discovery in genome graphs is finding the candidate regions of the graph that show evidence of dissimilarity from the sample’s reads.

### Finding candidate regions

The input required for finding candidate regions is a local graph, *n*, within the PanRG, the maximum likelihood path of both sequence and k-mers in *n, lmp*_*n*_ and *kmp*_*n*_ respectively, and a padding size *w* for the number of positions surrounding the candidate region to retrieve.

We define a candidate region, *r*, as an interval within *n* where coverage on *lmp*_*n*_ is less than a given threshold, *c*, for more than *l* and less than *m* consecutive positions. *m* acts to restrict the size of variants we are able to detect. If set too large, the following steps become much slower due to the combinatorial expansion of possible paths.

For a given read, *s*, that has a mapping to *r* we define *s*_*r*_ to be the subsequence of *s* that maps to *r*, including an extra *w* positions either side of the mapping. We define the pileup *P* _*r*_ as the set of all *s*_*r*_ ∈ *r*.

### Enumerating paths through candidate regions

For *r* ∈ *R*, where *R* is the set of all candidate regions, we construct a de Bruijn graph *G*_*r*_ from the pileup *P* _*r*_ using the GATB library(54). *A*_*L*_and *A*_*R*_are defined as sets of k-mers to the left and right of *r* in the local graph. They are anchors to allow re-insertion of new sequences found by *de novo* discovery into the local graph. If we cannot find an anchor on both sides, then we abandon *de novo* discovery for *r*. We use sets of k-mers for *A*_*L*_ and *A*_*R*_, rather than a single anchor k-mer, to provide redundancy in the case where sequencing errors cause the absence of some k-mers in *G*_*r*_. Once *G*_*r*_ is built, we define the start anchor k-mer, *a*_*L*_, as the first *a*_*L*_ ∈ *A*_*L*_ ∧ *a*_*L*_ ∈ *G*_*r*_. Likewise, we define the end anchor k-mer, *a*_*R*_, as the first *a*_*R*_ ∈ *A*_*R*_ ∧ *a*_*R*_ ∈ *G*_*r*_.

*T* _*r*_ is the spanning tree obtained by performing depth-first search (DFS) on *G*_*r*_, beginning from node *a*_*L*_. We define *p*_*r*_ as a path, from the root node *a*_*L*_ of *T* _*r*_ and ending at node *a*_*R*_, which fulfils the two conditions: 1) *p*_*r*_ is shorter than the maximum allowed path length. 2) No more than *k* nodes along *p*_*r*_ have coverage < *f* × *e*_*r*_, where *e*_*r*_ is the expected k-mer coverage for *r* and *f* is *n*_*r*_ × *s*, where *n*_*r*_ and *s* is a step size (0.1 by default).

*V* _*r*_ is the number of iterations of path enumeration for is the set of all *p*_*r*_. If |*V* _*r*_| is greater than a predefined threshold, then we have too many candidate paths, and we decide to filter more aggressively: *f* is incremented by *s –* effectively requiring more coverage for each *p*_*r*_ - and *V* _*r*_ discovery is abandoned for *r*.

### Pruning the path-space in a candidate region

As we operate on both accurate and error-prone sequencing reads, the number of valid paths in *G*_*r*_ can be very large. Primarily, this is due to cycles that can occur in *G*_*r*_ and exploring paths that will never reach our required end anchor *a*_*R*_. In order to reduce the path-space within *G*_*r*_ we prune paths based on multiple criteria. Critically, this pruning happens at each step of the graph walk (path-building).

We used a distance-based optimisation based on Rizzi et al (55). In addition to *T* _*r*_, obtained by performing DFS on *G*_*r*_, we produce a distance map *D*_*r*_ that results from running reversed breadth-first search (BFS) on *G*_*r*_, beginning from node *a*_*R*_. We say reversed BFS as we explore the predecessors of each node, rather than the successors. *D*_*r*_ is implemented as a binary search tree where each node in the tree represents a k-mer in *G*_*r*_ that is reachable from *a*_*R*_ via reversed BFS. Each node additionally has an integer attached to it that describes the distance from that node to *a*_*R*_.

We can use *D*_*r*_ to prune the path-space by 1) for each node *n* ∈ *p*_*r*_, we require *n* ∈ *D*_*r*_ and 2) requiring *a*_*R*_ be reached from *n* in, at most, *i* nodes, where *i* is defined as the maximum allowed path length minus the number of nodes walked to reach *n*.

If one of these conditions is not met, we abandon *p*_*r*_. The advantage of this pruning process is that we never explore paths that will not reach our required endpoint within the maximum allowed path length and when caught in a cycle, we abandon the path once we have made too many iterations around the cycle.

### Graph-based genotyping and optimal reference construction for multi-genome comparison

We use graph-based genotyping to output a comparison of samples in a VCF. A path through the graph is selected to be the reference sequence, and graph variation is described with respect to this reference. The chromosome field then details the local graph and the position field gives the position within the chosen reference sequence for possible variant alleles. The reference path for each local graph is chosen to be maximally close to the set of sample mosaic paths. This is achieved by reusing the mosaic path finding algorithm detailed in Supplementary Algorithm 3 on a copy of the k-mer graph with coverages incremented along each sample mosaic path, and a modified probability function defined such that the probability of a node is proportional to the number of samples covering it. This results in an optimal path, which is used as the VCF reference for the multi-sample VCF file.

For each sample and site in the VCF file, the mean forward and reverse coverage on k-mers tiling alleles is calculated. A likelihood is then independently calculated for each allele based on a Poisson model. An allele *A* in a site is called if: 1) *A* is on the sample mosaic path (i.e. it is on the maximum likelihood path for that sample); 2) *A* is the most likely allele to be called based on the previous Poisson model. Every allele not in the sample mosaic path will not satisfy 1) and will thus not be called. In the uncommon event where an allele satisfies 1), but not 2), we have an incompatibility between the global and the local choices, and then the site is genotyped as null.

### Comparison of variant-callers on a diverse set of *E. coli*

#### Sample selection

We used a set of 20 diverse *E. coli* samples for which matched Nanopore and Illumina data and a high-quality assembly were available. These are distributed across 4 major phylogroups of *E. coli* as shown in Figure 4. Of these, 16 were isolated from clinical infections and rectal screening swabs in ICU patients in an Australian hospital(56). One is the reference strain CFT073 that was resequenced and assembled by the REHAB consortium(57). One is from an ST216 cardiac ward outbreak (identifier: H131800734); the Illumina data was previously obtained(58) and we did the Nanopore sequencing (see below). The two final samples were obtained from Public Health England: one is a Shiga-toxin encoding *E. coli* (we used the identifier O63)(59), and the other an enteroaggregative *E. coli* (we used the identifier ST38)(60). Coverage data for these samples can be found in Supplementary Table 1.

#### PanRG construction

MSAs for gene clusters curated with PanX(27) from 350 RefSeq assemblies were downloaded from http://pangenome.de on 3rd May 2018. MSAs for intergenic region clusters based on 228 *E. coli* ST131 genome sequences were previously generated with Piggy(28) for their publication. The PanRG was built using *make_prg*. Two loci (GC00000027_2 and GC00004221) out of 37,428 were excluded because the combination of Clustal Omega and *make_prg* did not complete in reasonable time (∼24 hours) once *de novo* variants were added.

#### Nanopore sequencing of sample H131800734

DNA was extracted using a Blood & Cell Culture DNA Midi Kit (Qiagen, Germany) and prepared for Nanopore sequencing using kits EXP-NBD103 and SQK-LSK108. Sequencing was performed on a MinION Mk1 Shield device using a FLO-MIN106 R9.4 Spoton flowcell and MinKNOW version 1.7.3, for 48 hours.

#### Nanopore basecalling

Recent improvements to the accuracy of Nanopore reads have been largely driven by improvements in basecalling algorithms(61). All Nanopore data was basecalled with the methylation-aware, high-accuracy model provided with the proprietary guppy basecaller (version 3.4.5). In addition, 4 samples were basecalled with the default (methylation unaware) model for comparison (see Figure 5). Demultiplexing of the subsequent basecalled data was performed using the same version of the guppy software suite with barcode kits EXP-NBD104 and EXP-NBD114 and an option to trim the barcodes from the output.

#### Phylogenetic tree construction

Chromosomes were aligned using *MAFFT*(62) *v7.467 as implemented in Parsnp*(63) *v1.5.3. Gubbins* v2.4.1 was used to filter for recombination (default settings) and phylogenetic construction was carried out using *RAxML*(64) *v8.2.12 (GTR + GAMMA substitution model, as implemented in Gubbins*(65)).

#### Reference selection for mapping-based callers

A set of references was chosen for testing single-reference variant callers using two standard approaches, as follows. First, a phylogeny was built containing our 20 samples and 243 reference genomes from RefSeq. Then, for each of our 20 samples, the nearest RefSeq *E. coli* reference was found using Mash(66). Second, for each of the 20 samples, the nearest RefSeq reference in the phylogeny was manually selected; sometimes one RefSeq assembly was the closest to more than one of the 20. At an earlier stage of the project there had been another sample (making a total of 21) in phylogroup B1; this was discarded when it failed quality filters (data not shown). Despite this, the *Mash*/manual selected reference genomes were left in the set of mapping references, to evaluate the impact of mapping to a reference in a different phylogroup to all 20 of our samples.

#### Construction of truth assemblies

16/20 samples were obtained with matched Illumina and Nanopore data and a hybrid assembly. Sample H131800734 was assembled using the hybrid assembler *Unicycler*(67) with PacBio and Illumina reads followed by polishing with the PacBio reads using *Racon*(68), and finally with Illumina reads using *Pilon*(69). *A small 1kb artifactual contig was removed from the H131800734 assembly due to low quality and coverage*.

In all cases we mapped the Illumina data to the assembly, and masked all positions where the pileup of Illumina reads did not support the assembly.

#### Construction of a comprehensive and filtered truth set of pairwise SNPs

All pairwise comparisons of the 20 truth assemblies were performed *with varifier* (https://github.com/iqbal-lab-org/varifier), using subcommand *make_truth_vcf*. In summary, *varifier* compares two given genomes (referenced as G1 and G2) twice - first using *dnadiff*(70) and then using *minimap2/paftools*(53). The two output sets of pairwise SNPs are then joined and filtered. We create one sequence probe for each allele (a sequence composed of the allele and 50 bases of flank on either side taken from G1) and then map both to G2 using *minimap2*. We then evaluate these mappings to verify if the variant found is indeed correct (TP) or not (FP) as follows. If the mapping quality is zero, the variant is discarded to avoid paralogs/duplicates/repeats that are inherently hard to assess. We then check for mismatches in the allele after mapping and confirm that the called allele is the better match.

### Constructing a set of ground truth pan-genome variants

When seeking to construct a truth set of all variants within a set of bacterial genomes, there is no universal coordinate system. We start by taking all pairs of genomes and finding the variants between them, and then need to deduplicate them - e.g. when a variant between genomes 1 and 2 is the same as a variant between genomes 3 and 4, they should be identified; we define “the same” in terms of genome, coordinate and allele. An allele *A* in a position *P*_*A*_ of a chromosome *C*_*A*_ in a genome *G*_*A*_ is defined as a triple *A* = (*G*_*A*_, *C*_*A*_, *P*_*A*_). A pairwise variant *PwV* = {*A*_1_, *A*_2_} is defined as a pair of alleles that describes a variant between two genomes, and a pan-genome variant *PgV* = {*A*_1_, *A*_2_,…, *A*_*n*_} is defined as a set of two or more alleles that describes the same variant between two or more genomes. A pan-genome variant *P gV* can also be defined as a set of pairwise variants *PgV* = {*PwV* _1_, *PwV* _2_,…, *PwV* _*n*_}, as we can infer the set of alleles of *P gV* from the pairs of alleles in all these pairwise variants. Note that pan-genome variants are thus able to represent rare and core variants. Given a set of pairwise variants, we seek a set of pan-genome variants satisfying the following properties:

1. [Surjection]:
  a. each pairwise variant is in exactly one pan-genome variant;
  b. a pan-genome variant contains at least one pairwise variant;
2. [Transitivity]: if two pairwise variants *PwV* _1_ and *PwV* _2_ share an allele, then *PwV* _1_ and *PwV* _2_ are in the same pan-genome variant *P gV*;

We model the above problem as a graph problem. We represent each pairwise variant as a node in an undirected graph *G*. There is an edge between two nodes *n*_1_ and *n*_2_ if *n*_1_ and *n*_2_ share an allele. Each component (maximal connected subgraph) of *G* then defines a pan-genome variant, built from the set of pairwise variants in the component, satisfying all the properties previously described. Therefore, the set of components of *G* defines the set of pan-genome variants *P*. However, a pan-genome variant in *P* could: i) have more than one allele stemming from a single genome, due to a duplication/repeat; ii) represent biallelic, triallelic or tetrallelic SNPs/indels. For this evaluation, we chose to have a smaller, but more reliable set of pan-genome variants, and thus we filtered *P* by restricting it to the set of pan-genome variants *P* ′ defined by the variants *PgV* ∈ *P* such that: i) *P gV* has at most one allele stemming from each genome; ii) *PgV* is a biallelic SNP. *P* ′ is the set of 618,305 ground truth filtered pan-genome variants that we extracted by comparing and deduplicating the pairwise variants present in our 20 samples, and that we use to evaluate the recall of all the tools in this paper. Supplementary Figure 11 shows an example summarising the described process of building pan-genome variants from a set of pairwise variants.

### Subsampling read data and running all tools

All read data was randomly subsampled to 100x coverage using *rasusa* - the pipeline is available at https://github.com/iqbal-lab-org/subsampler. A *snakemake*(71) *pipeline to run the pandora* workflow with and without *de novo* discovery (see Figure 2d) is available at https://github.com/iqbal-lab-org/pandora_workflow. A *snakemake* pipeline to run *snippy, SAMtools, nanopolish* and *medaka* on all pairwise combinations of 20 samples and 24 references is available at https://github.com/iqbal-lab-org/variant_callers_pipeline.

### Evaluating VCF files

#### Calculating precision

Given a variant/VCF call made by any of the evaluated tools, where the input were reads from a sample (or several samples, in the case of *pandora*) and a reference sequence (or a PanRG, in the case of *pandora*), we perform the following steps to assess how correct a call is:

1. Construct a probe for the called allele, consisting of the sequence of the allele flanked by 150bp on both sides from the reference sequence. This reference sequence is one of the 24 chosen references for *snippy, SAMtools, nanopolish* and *medaka*; or the multi-sample inferred VCF reference for *pandora*;
2. Map the probe to the sample sequence using *BWA-MEM*(72);
3. Remove multi-mappings by looking at the Mapping Quality (MAPQ) measure(30) of the SAM records. If the probe is mapped uniquely, then its mapping passes the filter. If there are multiple mappings for the probe, we select the mapping *m*_1_ with the highest MAPQ if the difference between its MAPQ and the second highest MAPQ exceeds 10. If *m*_1_ does not exist, then there are at least two mappings with the same MAPQ, and it is ambiguous to choose which one to evaluate. In this case, we prefer to be conservative and filter this call (and all its related mappings) out of the evaluation;
4. We further remove calls mapping to masked regions of the sample sequence, in order to not evaluate calls lying on potentially misassembled regions;
5. Now we evaluate the mapping, giving the call a continuous precision score between 0 and 1. If the mapping does not cover the whole called allele, we give a score of 0. Otherwise, we look only at the alignment of the called allele (i.e. we ignore the flanking sequences alignment), and give a score of: number of matches / alignment length.

Finally, we compute the precision for the tool by summing the score of all evaluated calls and dividing by the number of evaluated calls. Note that here we evaluate all types of variants, including SNPs and indels.

#### Calculating recall

We perform the following steps to calculate the recall of a tool:

1. Apply the VCF calls to the associated reference using the VCF consensus builder (https://github.com/leoisl/vcf_consensus_builder), creating a mutated reference with the variants identified by the tool;
2. Build probes for each allele of each pan-genome variant previously computed (see Section “Constructing a set of ground truth pan-genome variants”);
3. Map all pan-genome variants’ probes to the mutated reference using *BWA-MEM*;
4. Evaluate each probe mapping, which is classified as a TP only if all bases of the allele were correctly mapped to the mutated reference. In the uncommon case where a probe multimaps, it is enough that one of the mappings are classified as TP;
5. Finally, as we now know for each pan-genome variant which of its alleles were found, we calculate both the pan-variant recall and the average allelic recall as per Section “*Pandora detects rare variation inaccessible to single-reference methods*”.

#### Filters

Given a VCF file with likelihoods for each genotype, the genotype confidence is defined as the log likelihood of the maximum likelihood genotype, minus the log likelihood of the next best genotype. Thus a confidence of zero means all alleles are equally likely, and high quality calls have higher confidences. In the recall/error rate plots of Figure 5 and Figures 6a,b, each point corresponds to the error rate and recall computed as previously described, on a genotype confidence (gt-conf) filtered VCF file with a specific threshold for minimum confidence.

We also show the same plot with further filters applied in Supplementary Figure 1. The filters were as follows. For Illumina data: for *pandora*, a minimum coverage filter of 5x, a strand bias filter of 0.05 (minimum 5% of reads on each strand), and a gaps filter of 0.8 were applied. The gaps filter means at least 20% the minimizer k-mers on the called allele must have coverage above 10% of the expected depth. As *snippy* has its own internal filtering, no filters were applied. For *SAMtools*, a minimum coverage filter of 5x was used. For Nanopore data: for *pandora*, a minimum coverage filter of 10x, a strand bias filter of 0.05, and a gaps filter of 0.6 were used. For *nanopolish*, we applied a coverage filter of 10x. We were unable to apply a minimum coverage filter to a *medaka* due to a software bug that prevents annotating the VCF file with coverage information.

#### Locus presence and distance evaluation

For all loci detected as present in at least one sample by *pandora*, we mapped the multi-sample inferred reference to all 20 sample assemblies and 24 references, to identify their true locations. To be confident of these locations, we employed a strict mapping using *bowtie2*(73) and requiring end-to-end alignments. From the mapping of all loci to all samples, we computed a truth locus presence-absence matrix, and compared it with *pandora*’s locus presence-absence matrix, classifying each *pandora* locus call as true/false positive/negative. Supplementary Figure 3 shows these classifications split by locus length. Having the location of all loci in all the 20 sample assemblies and the 24 references, we then computed the edit distance between them.

## Declarations

### Ethics approval and consent to participate

Not applicable

### Consent for publication

Not applicable

## Availability of data and materials

### Reproducibility

All input data for our analyses, including PanX’s and Piggy’s MSAs, PanRG, reference sequences, and sample data are publicly available (see Section “*Data availability*”). *Pandora*’s code, as well as all code needed to reproduce this analysis are also publicly available (see Section “*Code availability*”). Software environment reproducibility is achieved using Python virtual environments if all dependencies and source code are in Python, and using Docker(74) containers run with Singularity(75) otherwise. The exact commit/version of all repositories used to obtain the results in this paper can be retrieved with the git branch or tag *pandora_paper_tag1*.

### Data availability

- Gene MSAs from PanX, and intergenic MSAs from Piggy: doi.org/10.6084/m9.figshare.13204163;
- *E. Coli* PanRG: doi.org/10.6084/m9.figshare.13204172;
- Accession identifiers or Figshare links for the sample and reference assemblies, and Illumina and Nanopore reads are listed in Section D of the Supplementary file;
- Input packages containing all data to reproduce both the 4- and 20-way analyses described in the Results section are also available in Section D of the Supplementary file.

### Code availability

- *make_prg* (RCC graph construction algorithm): https://github.com/rmcolq/make_prg
- *pandora*: https://github.com/rmcolq/pandora
- *varifier*: https://github.com/iqbal-lab-org/varifier
- Pangenome variations pipeline taking a set of assemblies and returning a set of filtered pan-genome variants: https://github.com/iqbal-lab-org/pangenome_variations
- *pandora* workflow: https://github.com/iqbal-lab-org/pandora_workflow
- Run *snippy, samtools, nanopolish* and *medaka* pipeline: https://github.com/iqbal-lab-org/variant_callers_pipeline
- 4- and 20-way evaluation pipeline (recall/error rate curves etc):https://github.com/iqbal-lab-org/pandora_paper_roc
- Locus presence and distance from reference pipeline:https://github.com/iqbal-lab-org/pandora_gene_distance
- A master repository to reproduce everything in this paper, marshalling all of the above: https://github.com/iqbal-lab-org/paper_pandora2020_analyses

Although all containers are hosted on https://hub.docker.com/ (for details, see https://github.com/iqbal-lab-org/paper_pandora2020_analyses/blob/master/scripts/pull_containers/pull_containers.sh), and are downloaded automatically during the pipelines’ execution, we also provide Singularity(75) containers (converted from Docker containers) at doi.org/10.6084/m9.figshare.13204169.

Frozen packages with all the code repositories for *pandora* and the analysis framework can be found at doi.org/10.6084/m9.figshare.13204214.

## Authors’ contributions

RMC designed and implemented the fundamental data structures, and RCC, map and compare algorithms. MBH designed and implemented the *de novo* variant discovery component. LL optimised the codebase. RMC, MBH, LL designed and implemented (several iterations of) the evaluation pipeline, one component of which was written by MH. BL reimplemented and improved the RCC codebase. JH,SG,LP sequenced 18/20 of the samples. LWR, MBH, LL, KM, ZI analysed and visualised the 20-way data. ZI designed the study. ZI and RMC wrote the bulk of the paper, LL and MBH wrote sections, and all authors read and improved drafts.

## Acknowledgements

We are grateful to the REHAB consortium (https://modmedmicro.nsms.ox.ac.uk/rehab/) and the Transmission of Carbapenemase-producing Enterobacteriaceae (TRACE) study investigators for sharing sequencing data (for CFT073 and H131800734) in support of this work. We would like to thank Sion Bayliss and Ed Thorpe for discussions and help with Piggy. We are grateful to Kelly Wyres for sharing sequence data for the Australian samples, and to Tim Dallman and David Greig for sharing their data from Public Health England. We would like to thank the following for helpful conversations during the prolonged genesis of this project: Gil McVean, Derrick Crook, Eduardo Rocha, Bill Hanage, Ed Feil, Sion Bayliss, Ed Thorpe, Richard Neher, Camille Marchet, Rayan Chikhi, Kat Holt, Claire Gorrie, Rob Patro, Fatemeh Almodaresi, Nicole Stoesser, Liam Shaw, Phelim Bradley, and Sorina Maciuca.

