## Supplementary Material for "Nucleotide-resolution bacterial pan-genomics with reference graphs"

#### A. Supplementary figures

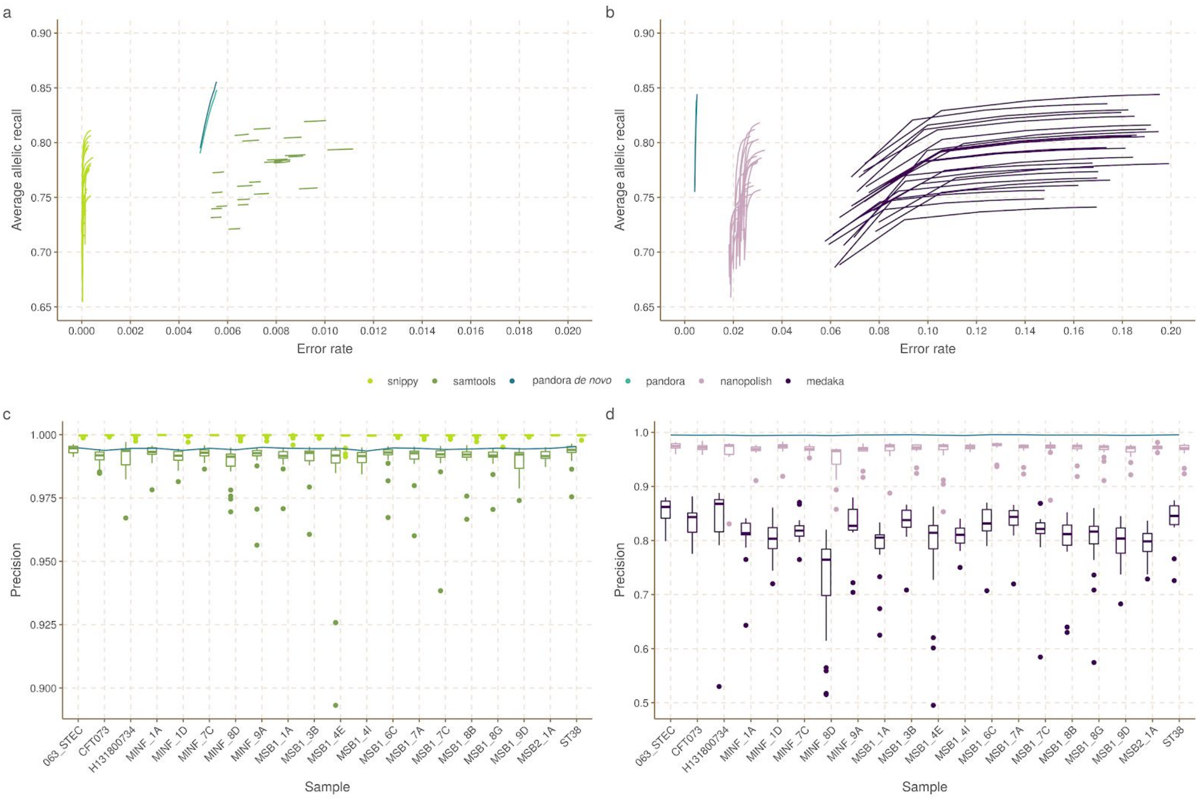


**Figure 1. Average allelic recall and error rate for *pandora* and other tools, with filters, on the 20-way dataset**. *a) The Average Allelic Recall and Error Rate curve for pandora, samtools and snippy on 100x of Illumina data; b) The Average Allelic Recall and Error Rate curve for pandora, medaka and nanopolish on 100x of Nanopore data; c) The precision (1 - error rate) of pandora, samtools and snippy on 100x of Illumina data. For samtools and snippy this varies with which of the 24 references was used; we display box plots showing the minimum, maximum and quartiles; the blue line connects pandora’s results; c) The precision of pandora (line plot), medaka and nanopolish (both box plots) on 100x of Nanopore data. Filters as follows. For Illumina data: for pandora, a coverage filter of 5x, strand bias 0.05, gaps filter 0.8 (explained in methods). As snippy has its own internal filtering, no filters were applied. For samtools, coverage filter 5x. For Nanopore data: for pandora coverage filter 10x, strand bias filter of 0.05 and a gaps filter of 0.6. For nanopolish, a coverage filter of 10x. For medaka, we were unable to apply the coverage filter due to a medaka software bug when annotating the VCF with coverage information.*

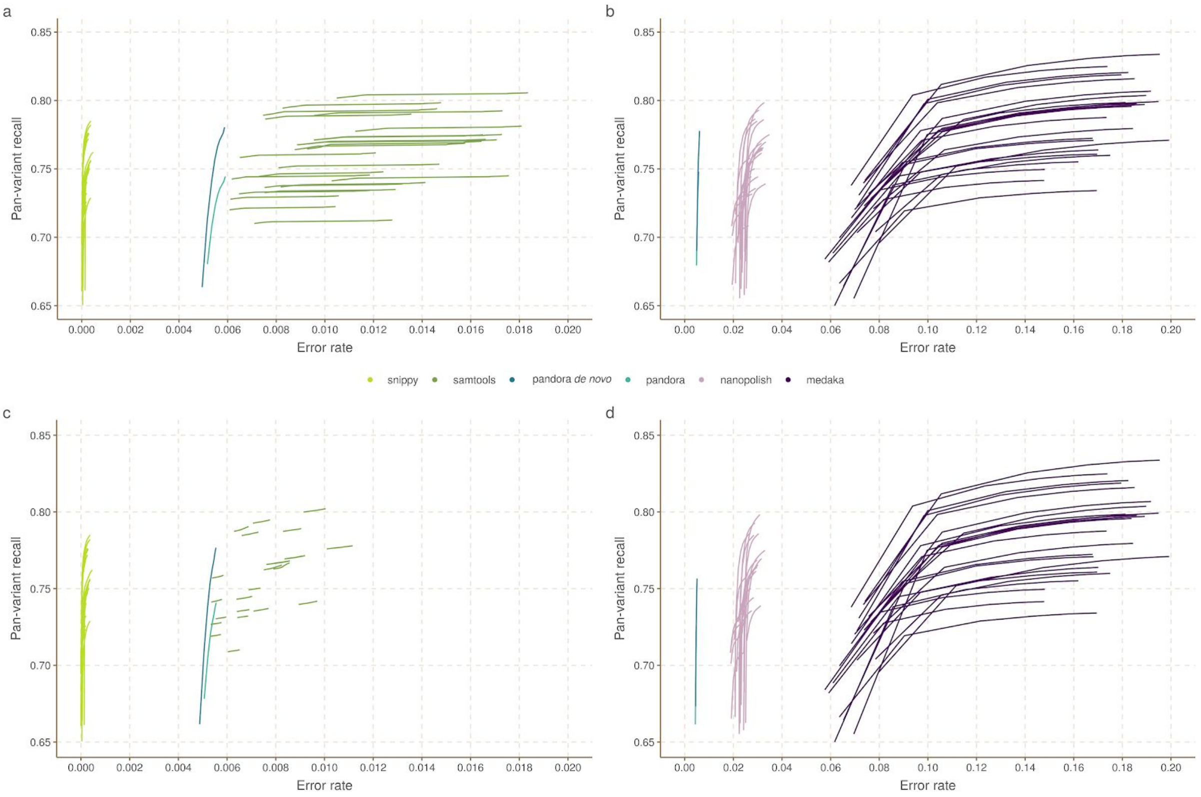


**Figure 2. Pan-variant recall and error rate for *pandora* and other tools, with and without filters, on the 20-way dataset.** *a) The Pan-variant Recall and Error Rate curve for pandora, samtools and snippy on 100x of Illumina data; b) The Pan-variant Recall and Error Rate curve for pandora, medaka and nanopolish on 100x of Nanopore data; c) Same as a), but with filters applied on the tools (for filters details, see caption of Supplementary Figure 1); d) Same as b), but with filters applied.*

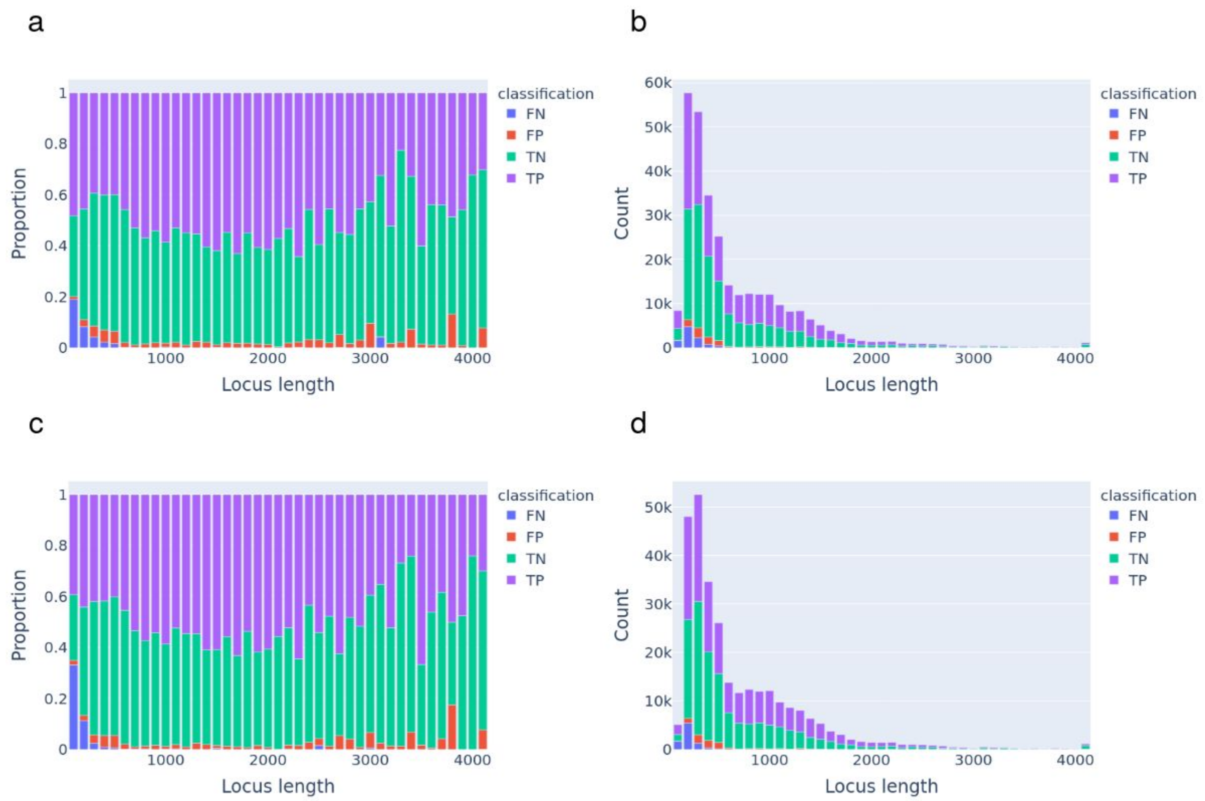


**Figure 3. Evaluation of pandora presence/absence classification of loci compared with truth.** *When pandora correctly identifies a locus as present or absent, these are labelled as true positive (TP)/true negative (TN) respectively. Similarly, wrongly identifying a gene as present or absent are labelled as false positive (FP)/false negative (FN). We display the counts and proportions of TP/TN/FP/FN loci broken down into bins according to their length. (a) On Illumina data, proportion of loci which are TN/TP/FN/FP; (b) As (a) but raw counts rather than proportion; (c) On Nanopore data, proportion of loci of a given length which are TN/TP/FN/FP; (d) As (c) but raw counts rather than proportion.*

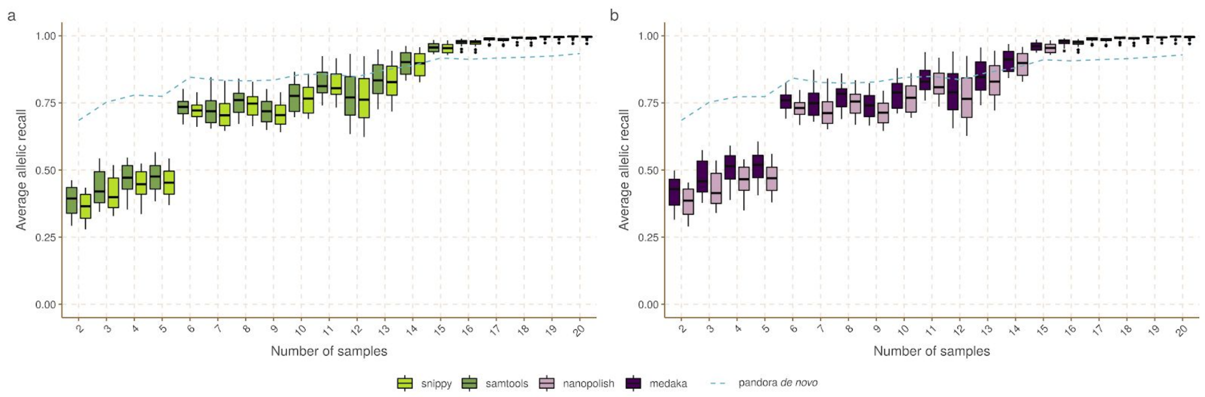


**Figure 4. Average allelic recall across the locus frequency spectrum.** *This figure shows the variation of AvgAR (average of the proportion of alleles of each pan-variant that are found, y-axis) across the locus frequency spectrum (number of samples in which the locus is present, x-axis) for pandora, snippy and samtools with Illumina data (a), and pandora, nanopolish and medaka with Nanopore data (b). For the reference-based tools, AvgAR depends on which reference is used, so box plots show the minimum, maximum and quartiles for samtools, snippy, nanopolish, and medaka; the blue line connects pandora results.*

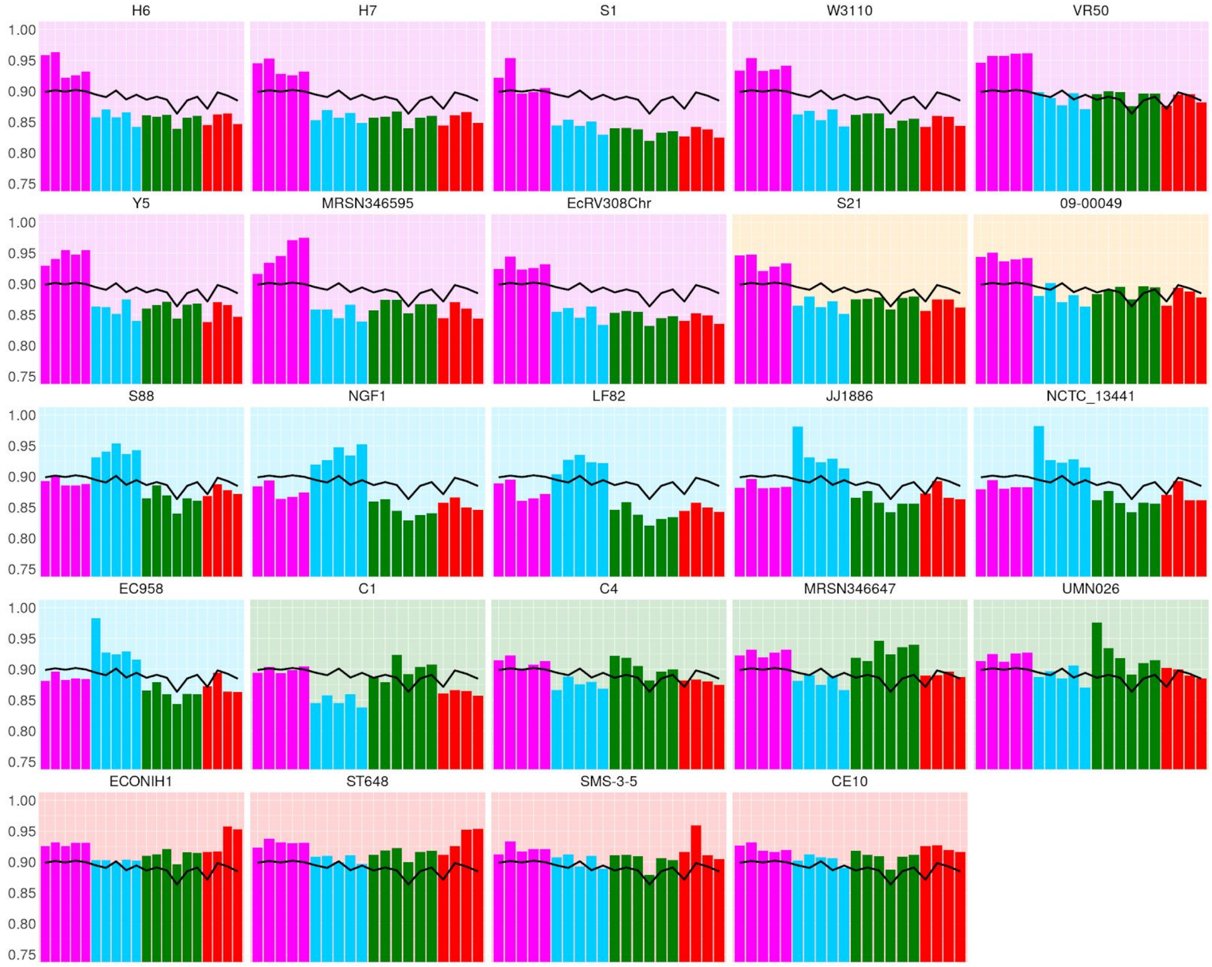


**Figure 5. Histograms showing how the phylogroup of a reference genome affects *snippy*'s recall in samples of various phylogroups**. *We show pandora recall (black line) and snippy recall (coloured bars) on the 20 samples; each histogram corresponds to the use of one of 24 references. The background colour indicates the reference's phylogroup (see Figure 4 inset); note that phylogroup B1 (yellow background) is an outgroup, containing no samples in this dataset.*

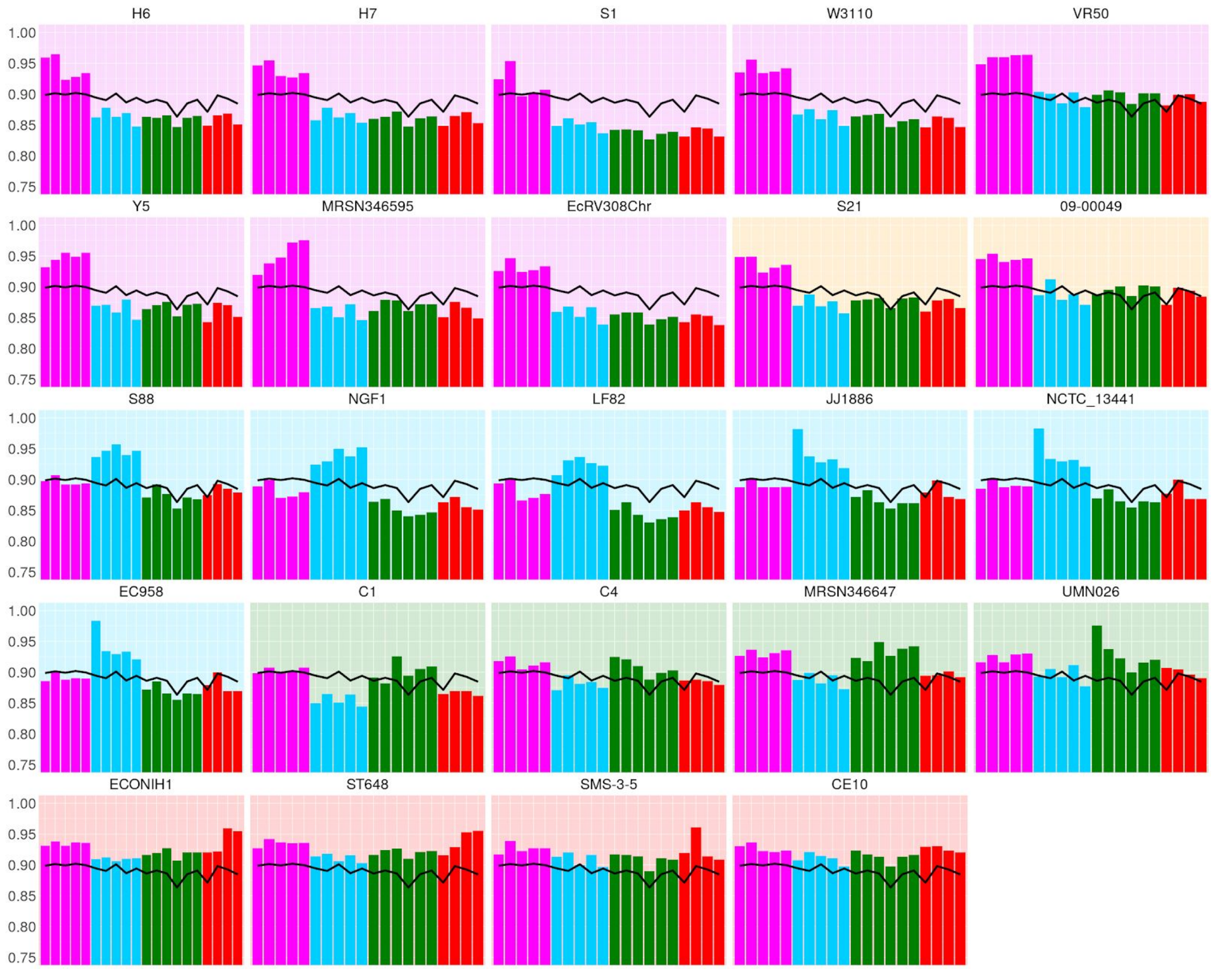


**Figure 6. Histograms showing how the phylogroup of a reference genome affects *samtools*' recall in samples of various phylogroups**. *We show pandora recall (black line) and samtools recall (coloured bars) on the 20 samples; each histogram corresponds to the use of one of 24 references. The background colour indicates the reference's phylogroup (see Figure 4 inset); note that phylogroup B1 (yellow background) is an outgroup, containing no samples in this dataset.*

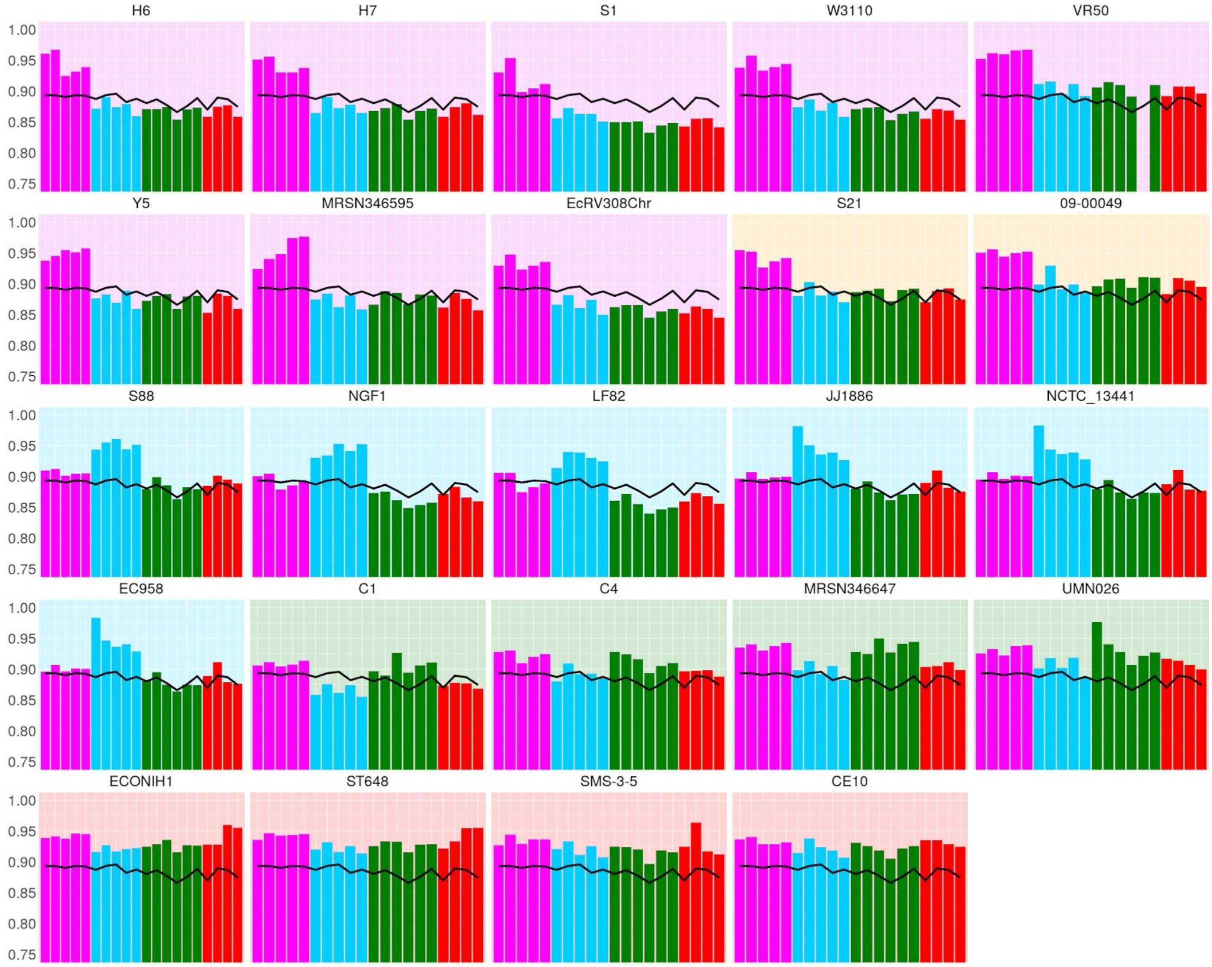


**Figure 7. Histograms showing how the phylogroup of a reference genome affects *medaka*'s recall in samples of various phylogroups**. *We show pandora recall (black line) and medaka recall (coloured bars) on the 20 samples; each histogram corresponds to the use of one of 24 references. The background colour indicates the reference's phylogroup (see Figure 4 inset); note that phylogroup B1 (yellow background) is an outgroup, containing no samples in this dataset.*

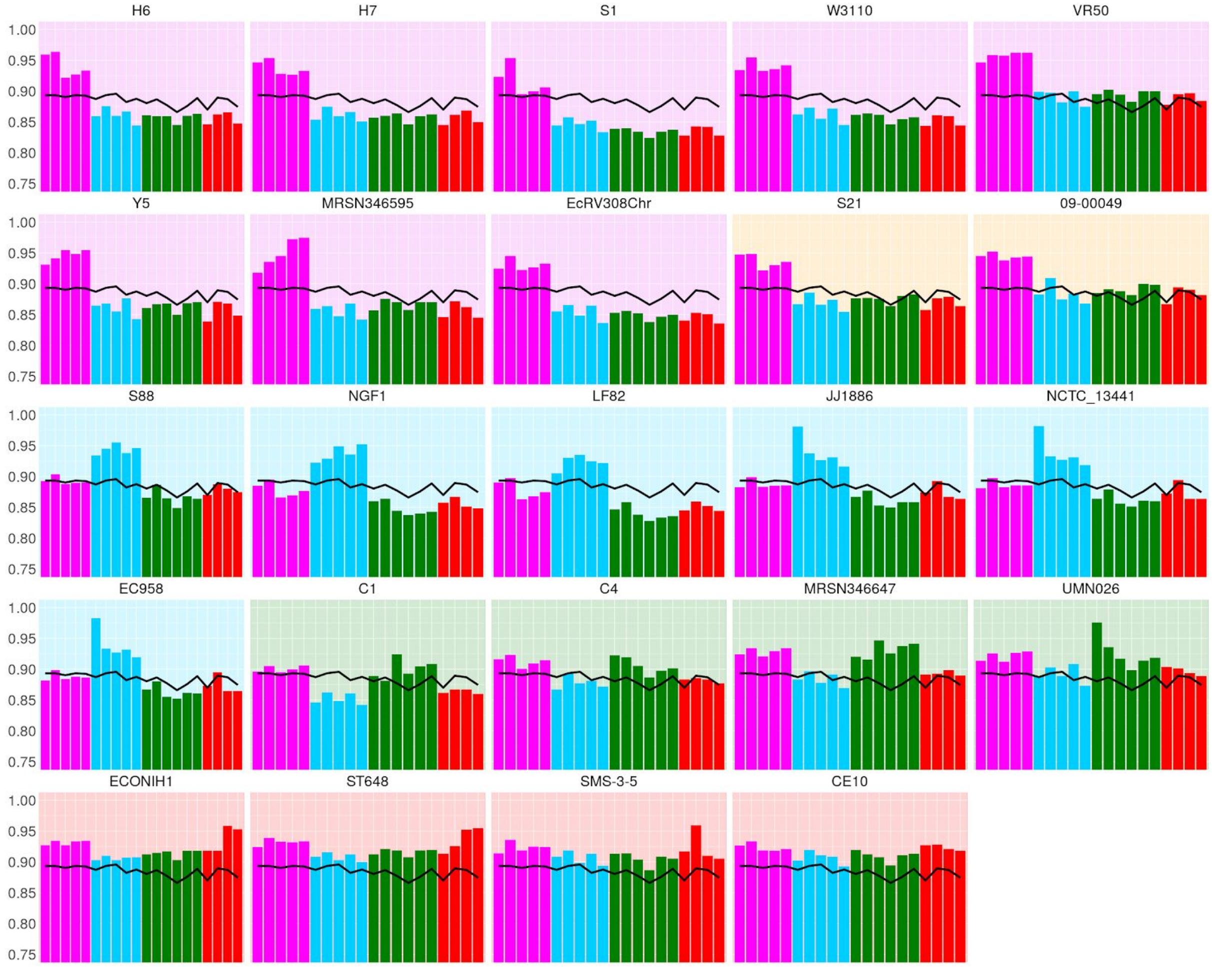


**Figure 8. Histograms showing how the phylogroup of a reference genome affects *nanopolish*'s recall in samples of various phylogroups**. *We show pandora recall (black line) and nanopolish recall (coloured bars) on the 20 samples; each histogram corresponds to the use of one of 24 references. The background colour indicates the reference's phylogroup (see Figure 4 inset); note that phylogroup B1 (yellow background) is an outgroup, containing no samples in this dataset.*

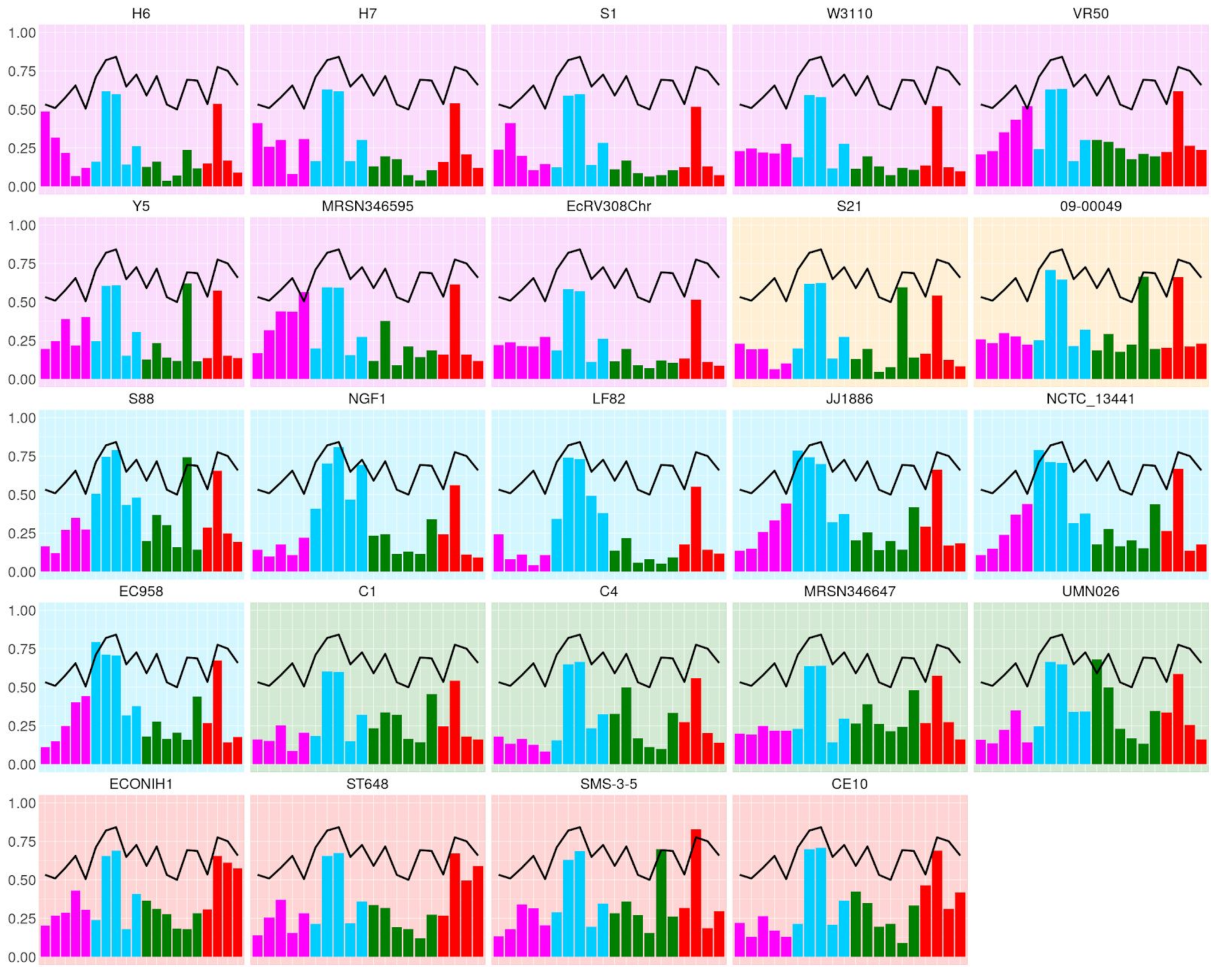


**Figure 9. Histograms showing how the phylogroup of a reference genome affects *snippy*'s recall of 2-variants in samples of various phylogroups**. *Restricting to variants present in precisely 2 of the 20 sample genomes, we show pandora recall (black line) and snippy recall (coloured bars) on the 20 samples; each histogram corresponds to the use of one of 24 references. The background colour indicates the reference's phylogroup (see Figure 4 inset); note that phylogroup B1 (yellow background) is an outgroup, containing no samples in this dataset.*

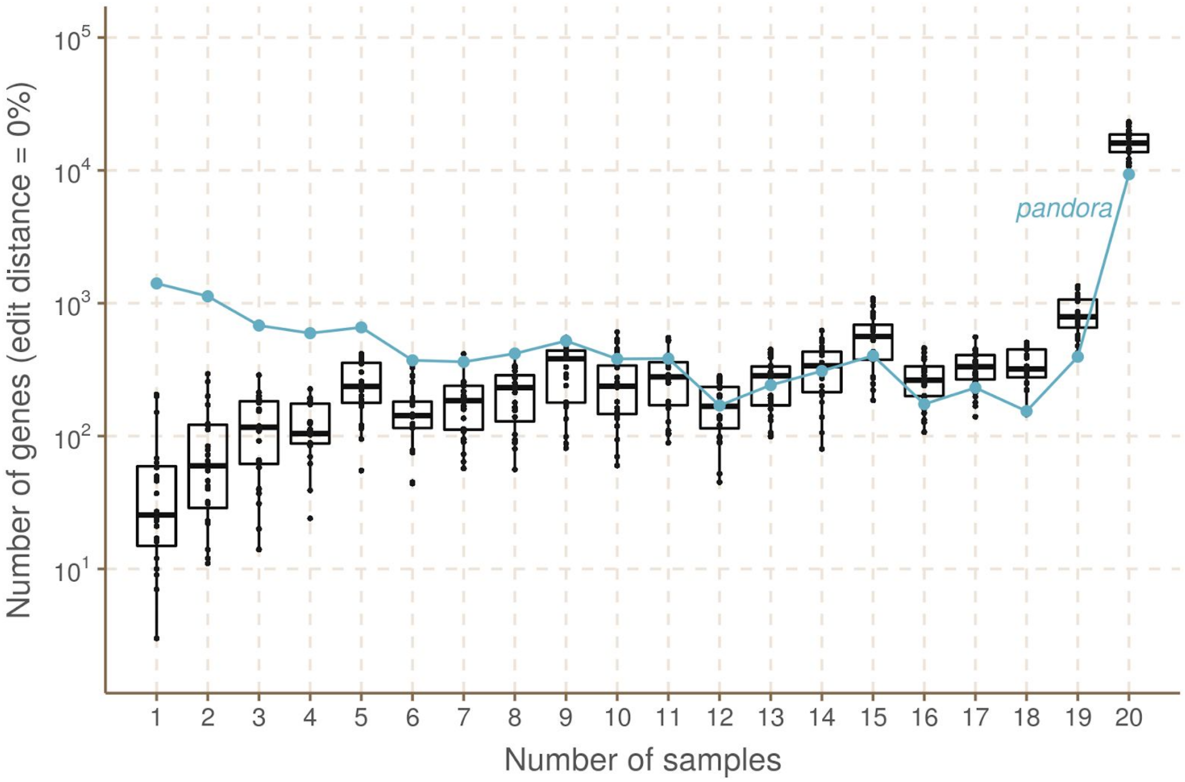


**Figure 10. How often do references have exactly the same sequence for a locus as a sample?** *We plot, on the y axis (log scale), the count of (locus, sample) pairs where the sample and reference have identical sequence for the locus (edit distance = 0). At each x coordinate we plot one point for each of the 20 reference genomes, with a box plot showing quartiles, minimum and maximum. The line plot shows results for the VCF-reference inferred by pandora.*

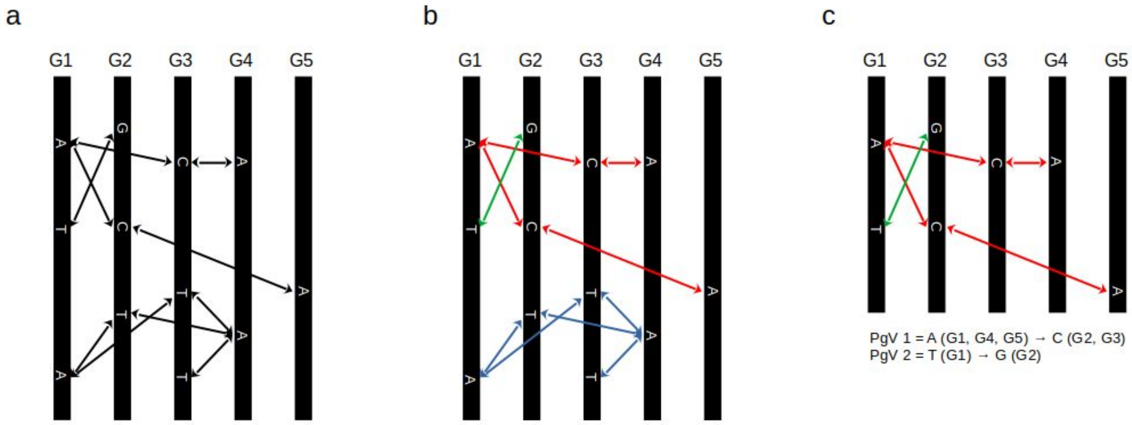


**Figure 11. Building pan-genome variants from a set of pairwise variants.** *a) Given five genomes, G1 to G5, the bi-directional arrows denote pairwise variants found by comparing pairs of genomes, and the letters denote the alleles. Note that pairwise genome comparison might miss real variants (FNs). For example, in this figure, we could have a SNP A (G5) → C (G3), which was not found by the pairwise comparison tool; b) Pairwise variants are coloured according to which pan-genome variants they belong to. This can be done through the transitivity property (see Methods in the main text); c) The blue pan-genome variant is filtered out because it has two alleles stemming from G3, probably due to a sequence duplication event. The two pan-genome variants we have in the end are: PgV 1 = A (G1, G4, G5) → C (G2, G3), a core variant present in all five genomes; and PgV 2 = T (G1) → G (G2), a rare variant present in only two genomes.*

#### B. Supplementary tables

| Sample | Illumina coverage | Nanopore coverage |
|---|---|---|
| O63_STEC | 100x | 100x |
| CFT073 | 94x | 100x |
| MINF_1A | 100x | 100x |
| MINF_1D | 100x | 100x |
| MINF_7C | 100x | 69x |
| MINF_8D | 100x | 100x |
| MINF_9A | 100x | 100x |
| MSB1_1A | 100x | 100x |
| MSB1_3B | 100x | 100x |
| MSB1_4E | 100x | 65x |
| MSB1_4I | 100x | 54x |
| MSB1_6C | 100x | 44x |
| MSB1_7A | 100x | 100x |
| MSB1_7C | 100x | 54x |
| MSB1_8B | 100x | 100x |
| MSB1_8G | 100x | 100x |
| MSB1_9D | 100x | 100x |
| MSB2_1A | 100x | 100x |
| H131800734 | 72x | 100x |
| ST38 | 53x | 100x |

**Table 1. Illumina and Nanopore read coverages of the *E. coli* samples used in the 20-way analysis.** *All read sets were downsampled to a maximum of 100x.*

| Tool | Reference | Technology | Error rate without/with filters | AvgAR without/with filters | Comment |
|---|---|---|---|---|---|
| Pandora no *de novo* | PanRG | Illumina | 0.59%/0.56% | 85%/84.8% | |
| Pandora with *de novo* | PanRG | Illumina | 0.59%/0.55% | 85.7%/85.6% | |
| Snippy | CP010170.1 | Illumina | 0.01%* | 73.7%* | Reference with best error rate |
| Snippy | NZ_CP009859.1 | Illumina | 0.04%* | 81.1%* | Reference with best AvgAR |
| Samtools | NZ_LM995446.1 | Illumina | 1.04%/0.57% | 73.3%/73.2% | Reference with best error rate |
| Samtools | NZ_CP009859.1 | Illumina | 1.84%/1% | 82.2%/82% | Reference with best AvgAR |
| Pandora no *de novo* | PanRG | Nanopore | 0.57%/0.48% | 85%/83.9% | |
| Pandora with *de novo* | PanRG | Nanopore | 0.61%/0.51% | 85.5%/84.4% | |
| Medaka | NZ_LM995446.1 | Nanopore | 14.79%* | 74.9%* | Reference with best error rate |
| Medaka | NZ_CP009859.1 | Nanopore | 19.54%* | 84.4%* | Reference with best AvgAR |
| Nanopolish | NC_007779.1 | Nanopore | 2.41%/2.26% | 73.9%/73.9% | Reference with best error rate |
| Nanopolish | NZ_CP009859.1 | Nanopore | 3.27%/3.09% | 81.8%/81.8% | Reference with best AvgAR |

* Filtered results not available.

**Table 2. Summary of AvgAR (average allelic recall) and error rate results for all tools in the 20-way analysis***: pandora (with and without de novo variant discovery), snippy, samtools, medaka and nanopolish (for these last four, we chose two references, that with the best AvgAR and that with the best error rate). Filtered results are also presented (for filters details, see caption of Supplementary Figure 1). In all cases the error rate quoted is for the point at the top-right of the recall/error rate curve (i.e. with no minimum genotype confidence filter), so for all tools the error rate could be improved further at a cost in recall.*

| Tool | Technology | Mean runtime | Max runtime | Nb of cores | Max RAM | Approximate input data size |
|---|---|---|---|---|---|---|
| Pandora | Illumina | 3h | 4.5h | 16 | 10.3GB | 8GB |
| Pandora | Nanopore | 4.1h | 5.9h | 16 | 10.7GB | 28GB |
| Snippy | Illumina | 0.1h | 0.2h | 4 | 3.2GB | 7GB |
| Samtools | Illumina | 0.3h | 0.5h | 1 | 1.1GB | 7GB |
| Medaka | Nanopore | 0.3h | 0.6h | 4 | 5GB | 27GB |
| Nanopolish | Nanopore | 4.6h | 12.3h | 16 | 13GB | 1.4TB |

**Table 3. Computational performance for single-sample analysis by each tool.** *We present, for each tool, the mean and the maximum runtime (wall clock time), number of cores used, maximum RAM usage and approximate input data size. For pandora these values refer to mapping and performing de novo variant discovery on each sample.*

#### C. Supplementary Algorithms

**ALGORITHM** Find the minimizing $k$-mers for a graph

**Input :** G – a local graph
**Output:** K – a $k$-mer-graph of minimizing $k$-mers

```
Function MinimizerSketch(G)
    L ← ∅
    T ← ∅
    X ← walk(0, 0, w, k)
    for x ∈ X do
        S ← minimize(x)
        K ← add(∅, S)
        L ← L ∪ S
    while L ≠ ∅ do
        m := (h, p, r) ← L.extract()
        if m == G.end() then
            T ← T ∪ {m}
            continue
        X ← {x \ m for x ∈ shift(m)}
        while X ≠ ∅ do
            x ← X.extract_last()
            y ← x.get_last()
            if min{φ(π(s_y), 0), φ(π(s_y), 1)} ≤ min{φ(π(s_p), 0), φ(π(s_p), 1)}
             then
                K ← add(m, y)
                L ← L ∪ {y}
            else if |x| == w then
                S ← minimize(x)
                K ← add(m, S)
                L ← L ∪ S
            else if y == G.end() then
                T ← T ∪ {m}
            else
                X ← X ∪ {shift(x)}
    K ← add(T, ∅)
    return K
```

**Algorithm 1. Minimizer sketch algorithm.**

**ALGORITHM** Quasi-map

**Input :** R – a read
$\mathcal{I}$ – the PanRG Index
$m$ – the maximum gap length (in bases) between $k$-mers in a cluster on a read
$t$ – the minimum number of hits in a cluster
**Output:** $\mathcal{C}$ – Clusters of hits corresponding to mapped regions

**Function** `QuasiMap`$(R, \mathcal{I})$
  M $\leftarrow$ `MinimizerSketch`(R)
  A $\leftarrow \emptyset$
  **for** $(h, p, r) \in M$ **do**
    **for** $(j, \mathbf{q}, r') \in \mathcal{I}[h]$ **do**
      **if** $r \equiv r'$ **then**
        Append $(j, 0, p, \mathbf{q})$ to A
      **else**
        Append $(j, 1, p, \mathbf{q})$ to A
  Sort A $= \{(j, r, p, \mathbf{q})\}$ by the 4 values of tuple
  $\mathcal{C} \leftarrow \emptyset$
  Cluster $\leftarrow \emptyset$
  $a_{prev} \leftarrow$ None
  **for** $a_{curr} = (j, r, p, \mathbf{q}) \in A$ **do**
    **if** $j_{curr} \neq j_{prev}$ ***or*** $r_{curr} \neq r_{prev}$ ***or*** $\|p_{curr} - p_{prev}\| > m$ **then**
      **if** *Cluster*.size() $> t$ **then**
        $\mathcal{C}$.append(Cluster)
      Cluster $\leftarrow \emptyset$
    Cluster.append($a_{curr}$)
    $a_{prev} = a_{curr}$
  **return** $\mathcal{C}$

**Algorithm 2. Quasi-mapping reads to the PanRG.**

**ALGORITHM** Find the mosaic path

```
Input  : K – a directed acyclic k-mer graph with n nodes
         t – terminus probability threshold
Output: p = {p_1, ..., p_m} – a path through K

Function FindMosaic (K)
    M ← arr(0, n)
    len ← arr(0, n)
    prev ← arr(n, n)
    for i = n, ..., 1 do
        max_mean ← − inf
        max_length ← 0
        for j ∈ outnodes(i) do
            if (j.is_terminus() and t > max_mean)
              or M[j]/len[j] > max_mean
              or (M[j]/len[j] ≡ max_mean and len[j] > max_len) then
                if j.is_terminus() then
                    max_mean = t
                else
                    max_mean = M[j]/len[j]
                    max_len = len[j]
                M[i] = M[j] + prob(i)
                len[i] = len[j] + 1
                prev[i] = j
    prev_node = prev[0]
    p ← ∅
    while prev_node < n do
        p.append(prev_node)
        prev_node = prev[prev_node]
    return p
```

**Algorithm 3. Mosaic sequence inference algorithm.**

### D. Data availability

#### Input packages

To facilitate the reproducibility of our pipelines, we provide input packages containing all data required to run both the 4- and 20-way analyses described in the Results section of the main text. All data can be found at this ftp site: ftp://ftp.ebi.ac.uk/pub/software/pandora/2020/

| Package | Link | Description |
|---|---|---|
| 4-way input (21 GB) | ftp://ftp.ebi.ac.uk/pub/software/pandora/2020/data_4_way_methylation/ | Contains the input data to run the 4-way analysis with Nanopore reads before and after guppy methylation-aware basecalling. |
| 20-way input - no fast5s (38 GB) | ftp://ftp.ebi.ac.uk/pub/software/pandora/2020/data_no_fast5s/ | Contains the input data to run the 20-way analysis, without the fast5 data. Nanopolish can't therefore be run, but this makes it a much lighter package. |
| 20-way input - full (1.3 TB) | ftp://ftp.ebi.ac.uk/pub/software/pandora/2020/data/ | Contains the complete input data to run the 20-way analysis, including the fast5 data. |

Details of the data inside this package (assemblies accessions, reads biosamples, etc) are presented in the next sections.

#### Sample assemblies

| Sample | Accession ID / Figshare link |
|---|---|
| O63_STEC | GCA_004804185.1 |
| CFT073 | https://ndownloader.figshare.com/files/16262078 |
| H131800734 | doi.org/10.6084/m9.figshare.13204184 |
| ST38 | doi.org/10.6084/m9.figshare.13204187 |
| MINF_1A | GCA_904863375 |
| MINF_1D | GCA_904863105 |
| MINF_7C | GCA_904863235 |

| | |
|---|---|
| MINF_8D | GCA_904866345 |
| MINF_9A | GCA_904863255 |
| MSB1_1A | GCA_904863385 |
| MSB1_3B | GCA_904863305 |
| MSB1_4E | GCA_904866455 |
| MSB1_4I | GCA_905071835.1 |
| MSB1_6C | GCA_904866495 |
| MSB1_7A | GCA_904864615 |
| MSB1_7C | GCA_904864555 |
| MSB1_8B | GCA_904865655 |
| MSB1_8G | GCA_904866225 |
| MSB1_9D | GCA_904863445 |
| MSB2_1A | GCA_904864635 |

All the masks for the assemblies are available at doi.org/10.6084/m9.figshare.13204175.

##### Reference assemblies

| Reference | Accession ID | Strain name (e.g. see Figure 4 in the main text) |
|---|---|---|
| CU928163.2 | GCA_000026325.1 | UMN026 |
| CP018206.1 | GCA_001890365.1 | MRSN346647 |
| CP010116.1 | GCA_001900295.1 | C1 |
| CP010121.1 | GCA_001900315.1 | C4 |
| CP010170.1 | GCA_001900435.1 | H6 |
| CP010171.1 | GCA_001900455.1 | H7 |
| CP010230.1 | GCA_001900925.1 | S21 |
| CP010226.1 | GCA_001901315.1 | S1 |
| NC_007779.1 | GCF_000010245.2 | W3110 |
| NC_010498.1 | GCF_000019645.1 | SMS-3-5 |
| NC_011742.1 | GCF_000026285.1 | S88 |

| | | |
|---|---|---|
| NC_017646.1 | GCF_000227625.1 | CE10 |
| NC_011993.1 | GCF_000284495.1 | LF82 |
| NZ_HG941718.1 | GCF_000285655.3 | EC958 |
| NC_022648.1 | GCF_000493755.1 | JJ1886 |
| NZ_CP009859.1 | GCF_000784925.1 | ECONIH1 |
| NZ_LM995446.1 | GCF_000952955.1 | EcRV308Chr |
| NZ_CP011134.1 | GCF_000968515.1 | VR50 |
| NZ_CP008697.1 | GCF_001485455.1 | ST648 |
| NZ_CP016007.1 | GCF_001660585.1 | NGF1 |
| NZ_CP015228.1 | GCF_001677495.1 | 09-00049 |
| NZ_CP013483.1 | GCF_001860505.1 | Y5 |
| NZ_CP018109.1 | GCF_001886755.1 | MRSN346595 |
| NZ_LT632320.1 | GCF_900119685.1 | NCTC_13441 |

##### Illumina reads

| Sample | Run ID |
|---|---|
| O63_STEC | SRR6144123 |
| CFT073 | SRR8482585 |
| H131800734 | SRR5367999 |
| ST38 | SRR5470155 |
| MINF_1A | ERR1015394 |
| MINF_1D | ERR1015416 |
| MINF_7C | ERR1015413 |
| MINF_8D | ERR1015419 |
| MINF_9A | ERR1015399 |
| MSB1_1A | ERR1023706 |
| MSB1_3B | ERR1023714 |
| MSB1_4E | ERR1015350 |

| | |
|---|---|
| MSB1_4I | ERR1015376 |
| MSB1_6C | ERR1015333 |
| MSB1_7A | ERR1023710 |
| MSB1_7C | ERR1015334 |
| MSB1_8B | ERR1015326 |
| MSB1_8G | ERR1015365 |
| MSB1_9D | ERR1015345 |
| MSB2_1A | ERR1015392 |

##### Nanopore reads

All original Nanopore reads were basecalled with the default (non methylation-aware) Guppy model. We used the original Nanopore reads only for four samples (O63_STEC, CFT073, H131800734 and ST38). The run accession IDs for the Nanopore reads for these four samples are shown in the table below. All the Nanopore reads for all the 20 samples were then re-basecalled using the methylation-aware Guppy model (dna_r9.4.1_450bps_modbases_dam-dcm-cpg_hac_prom.cfg), and are available as part of the 20-way input package. The fast5 data for each sample can also be found in the full 20-way input package.

| Sample | Run ID |
|---|---|
| O63_STEC | SRR8792695 |
| CFT073 | SRR8494940 |
| H131800734 | Available inside the 4-way input package. |
| ST38 | SRR6702264 |

#### E. Supplementary animations

doi.org/10.6084/m9.figshare.13204199
**Animation 1. Recursive clustering construction.**

doi.org/10.6084/m9.figshare.13204211
**Animation 2. *Pandora* vs *snippy* recall split by pan-genome variant frequency, and *snippy* reference.** *Each frame shows pandora recall (black line) and snippy recall (coloured bars) on the 20 samples, on pan-genome variants with a specific frequency. The background colour denotes the reference's phylogroup.*
